# The underlying mechanisms of alignment in error backpropagation through arbitrary weights

**DOI:** 10.1101/2021.06.12.447639

**Authors:** Alireza Rahmansetayesh, Ali Ghazizadeh, Farokh Marvasti

**Affiliations:** Electrical Engineering Department, Sharif University of Technology, Tehran, Iran

**Keywords:** backpropagation, feedback alignment, weight transport problem, bio-inspired artificial neural networks, weight normalization

## Abstract

Understanding the mechanisms by which plasticity in millions of synapses in the brain is orchestrated to achieve behavioral and cognitive goals is a fundamental quest in neuroscience. In this regard, insights from learning methods in artificial neural networks (ANNs) and in particular supervised learning using backpropagation (BP) seem inspiring. However, the implementation of BP requires exact matching between forward and backward weights, which is unrealistic given the known connectivity pattern in the brain (known as “weight transport problem”). Notably, it has been shown that under certain conditions, error <u>B</u>ack<u>P</u>ropagation <u>T</u>hrough <u>A</u>rbitrary <u>W</u>eights (BP-TAW) can lead to a partial alignment between forward and backward weights (weight alignment or WA). This learning algorithm, which is also known as feedback alignment (FA), can result in surprisingly good degrees of accuracy in simple classification tasks. However, the underlying mechanisms and mathematical basis of WA are not thoroughly understood. In this work, we show that the occurrence of WA is governed by statistical properties of the output and error signals of neurons, such as autocorrelation and cross-correlation, and can happen even in the absence of learning or reduction of the loss function. Moreover, we show that WA can be improved significantly by limiting the norm of input weights to neurons and that such a weight normalization (WN) method can improve the classification accuracy of BP-TAW. The findings presented can be used to further improve the performance of BP-TAW and open new ways for exploring possible learning mechanisms in biological neural networks without exact matching between forward and backward weights.

## 1 Introduction

For the past four decades, BP has been the dominant learning method used in artificial neural networks (Rumelhart et al., 1985). However, BP is known to be implausible in the nervous system (Stork, 1989; Crick, 1989; Song et al., 2020). One of its major issues is known as “weight transport problem” (Grossberg, 1987) which refers to the requirement that backward weights should be precisely matched with the forward weights so that accurate error signals are backpropagated to the early layers for efficient supervised learning. However, in the brain, axons transmit information unidirectionally, and to date, no explicit mechanism that guarantees a match between backward and forward weights is reported.

Despite differences in natural and artificial learning mechanisms, striking similarities between the activity of neurons in the brain and that of artificial ones trained by BP have been reported (Zipser and Andersen, 1988; Khaligh-Razavi and Kriegeskorte, 2014; Cadieu et al., 2014; Cichy et al., 2016; Nayebi et al., 2018), and possibilities for calculation of approximate gradient directions in the brain are suggested (Whittington and Bogacz, 2019, 2017; Lillicrap et al., 2020; Xie and Seung, 2003). In particular, it has been shown that learning occurs even without exact weight transport (Kolen and Pollack, 1994; Liao et al., 2016) and when arbitrary weights that are distinct from forward ones backpropagate error signals to early layers (Lillicrap et al., 2016), a method which we refer to as BP-TAW. During the learning process using BP-TAW, the angle between the forward weight matrices and the transpose of backward weight matrices in each layer reduces and this partial alignment leads to weight update directions that are partially aligned with weight update directions calculated by BP, thereby providing effective teaching signals (Lillicrap et al., 2016). It has also been shown that by a learning algorithm known as direct feedback alignment (DFA), learning can occur even when errors are passed directly from the output layer to each hidden layer through direct arbitrary backward weights (Nøkland, 2016; Refinetti et al., 2020; Frenkel et al., 2019; Launay et al., 2019; Baldi et al., 2018). Interestingly, these sub-optimal calculations of gradient directions (compared to BP) can lead to a surprisingly good degree of learning accuracy, comparable to BP in shallow networks and simple tasks but with a drop in accuracy in deep convolutional networks and complex tasks (Bartunov et al., 2018; Launay et al., 2019; Moskovitz et al., 2018).

Although there are some investigations on the favorable conditions for WA and for the improvement of learning using BP-TAW (Refinetti et al., 2020; Frenkel et al., 2019; Moskovitz et al., 2018; Akrout et al., 2019; Kunin et al., 2020; Xiao et al., 2018; Baldi et al., 2018; Lillicrap et al., 2016), the underlying mathematical basis for the success of BP-TAW in training ANNs is not fully understood. For instance, previous works have explored some of the conditions that lead to WA in the special case of linear networks (Lillicrap et al., 2016; Frenkel et al., 2019). It has been shown that if all forward weights are initialized to zero and the input and desired output of a network are kept constant during iterations, each backward weight matrix becomes a scalar multiple of the Moore-Penrose pseudo-inverse of forward weight matrices (Lillicrap et al., 2016), or a scalar multiple of the Moore-Penrose pseudo-inverse of the product of forward weight matrices in the case of DFA (Frenkel et al., 2019). Under these circumstances, update directions of BP-TAW are an approximation of the Gauss-Newton optimization method (Lillicrap et al., 2016). However, these proofs cannot explain the occurrence of WA for arbitrary initialization of weights (supplementary figure 13 of Lillicrap et al. 2016) and for nonlinear networks.

There are other preliminary attempts to explain the occurrence of alignment. In particular, Lillicrap et al. 2016 have provided an insight into the mechanics of FA by freezing forward weights in different stages of the learning process of an ANN trained by BP-TAW, showing that information about backward weight matrix of each layer (*B*_*ℓ*_ in Fig. 1) gradually accumulates in the earlier forward weight matrix (*W*_*ℓ* −1_ in Fig. 1) and then flows into the next forward weight (*W*_*ℓ*_ in Fig. 1) such that each forward weight matrix aligns with its corresponding backward weight matrix (*W*_*ℓ*_ and *B*_*ℓ*_ in Fig. 1). It has been also noted that under a restricted condition where input of a 2-layer linear network is white noise and the network is trained to learn a linear function, continuous growth of norm of weight matrices results in alignment (supplementary note 12 of Lillicrap et al. 2016). We will show that although the original form of BP-TAW is accompanied by a continuous growth of the norms of the weight matrices, this can be detrimental to WA and that limiting the norms of weights can improve WA.

**Figure 1.**
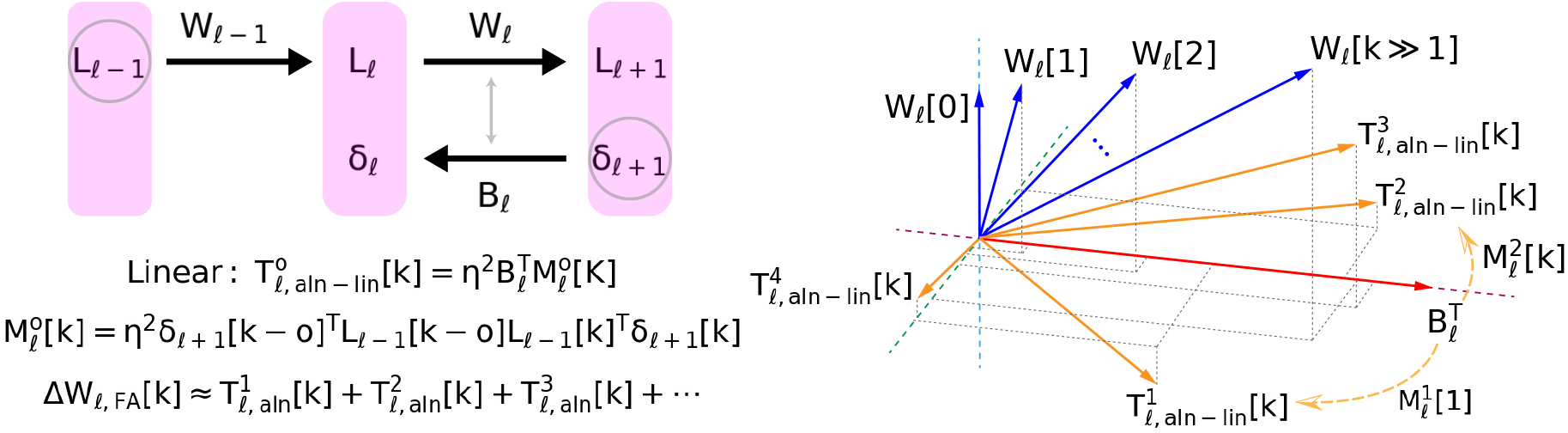
The underlying mechanism of weight alignment in BP-TAW. Expansion of Δ*W*_*ℓ, FA*_[*k*] reveals alignment terms 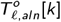 (equations 3, 4, and 5). In linear alignment terms, 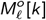 acts as a transformation matrix on 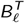 and if it partially preserves the direction of 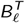 after the matrix multiplication, it propels *W*_*ℓ*_ [*k*] towards 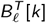. *L*_*ℓ*_[*k*] and *δ*_*ℓ*_ [*k*] denote the matrices of output and error signals of neurons, respectively, where each row of them corresponds to a data point of the mini-batch at the iteration *k* and each column of them corresponds to a neuron in the layer *ℓ*. Due to the structure of 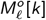, alignment terms can robustly propel forward weight matrices (*W*_*ℓ*_) towards transpose of fixed random backward weight matrices 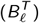 under a variety of conditions depending on neural activity. Note that this is a simplified diagram of underlying mechanism of WA. In practice, at each iteration *k* there are *k* alignment terms, and depending on the neural activity, each of them may or may not align with 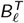. Moreover, in nonlinear ANNs, nonlinearity affects the structure of alignment terms to some extent (equation 4, see also Supplementary Note 2). An order (1 ≤ *o* ≤ *k*) is assigned to alignment terms since in each of them the activity of neurons in iterations of *k* and *k* − *o* (lag of *o*) are multiplied together.

In this work, we first explore the mathematical basis of WA in the general case. We show that occurrence of alignment does not rely on reduction of the loss function; rather, it is driven by statistical properties of neural activity such as cross-correlation and autocorrelation of error and output signals of neurons. Afterwards, we dissect WA in a practical application of BP-TAW for training a specific nonlinear 5-layer ANN on the MNIST dataset. We show that the relative similarity of data points that belong to a single category compared to the ones that belong to different categories contributes to alignment by shaping cross-correlated neural activity. Then, we will compare the trajectories of the forward weights of the network trained by BP and BP-TAW using low dimensional embedding and show that in BP-TAW the network convergences to a different and less optimal local minimum than BP, but the performance of BP-TAW can be improved by limiting the Frobenius norms of the input weights to each neuron.

## 2 Results

### 2.1 Driving alignment terms to explain the occurrence of alignment

Consider a conventional *d* layer ANN. We denote the matrices of forward weights (with arbitrary initialization), internal states of neurons, and output signals of neurons by 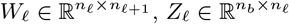, and *L*_*ℓ*_ = *f*(*Z*_*ℓ*_), respectively, where *n*_*b*_ is batch size, *n*_*ℓ*_ is the number of neurons in the layer *R* (network dimensions), and *f* (·) is an element-wise activation function (the following analysis still holds true if the batch size is variable among mini-batches; however, for simplicity, we assume that it is a constant number for all of them). For 0 *< R* ≤ *d*, internal states of neurons in the layer *R* are calculated according to *Z*_*ℓ*_ = *L*_*ℓ* −1_*W*_*ℓ* −1_ + **b**_*ℓ*_ where **b**_*ℓ*_ is the bias vector of the layer *R* and the addition of a matrix with a row vector is defined as adding the vector to each row of the matrix. We denote input, output, and desired output matrices of the network by 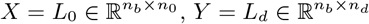, and 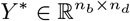, respectively.

In BP-TAW (Lillicrap et al., 2016), the error is backpropagated through constant arbitrary matrices (different from forward weights) denoted by 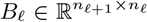, and weight update directions are calculated at each iteration *k* ≥ 0 according to

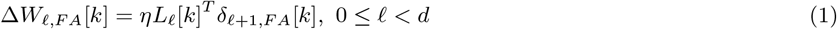

where *η* is the learning rate and error signals of neurons are

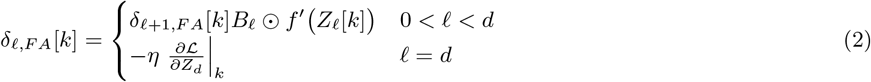

and ℒ (*Y, Y* ^*^) is the loss function and ⊙ denotes element-wise matrix multiplication (in the order of operations, it has less priority than matrix multiplication).

To demonstrate why this update rule leads to alignment, Δ*W*_*ℓ, F A*_[*k*] can be expanded by taking successive steps backward along the iterations and substituting every *W*_*ℓ*_ [*k* −*o*] for 0 ≤ *o < k*. Assuming update steps to be small, by applying first-order Taylor approximation we have

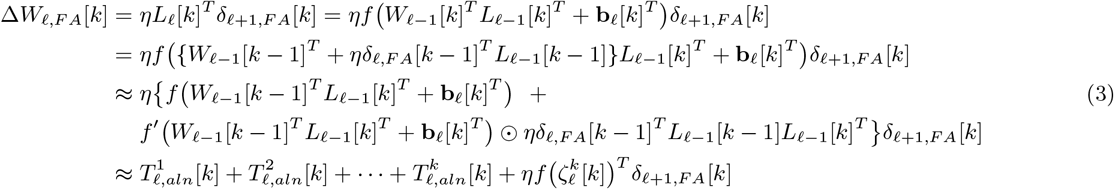

where 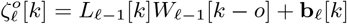 and for 1 ≤ *o* ≤ *k* and 0 *< ℓ < d* we define

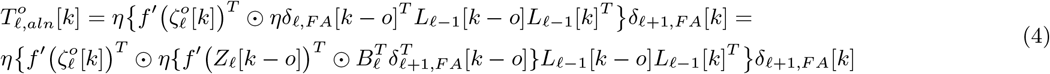

as *alignment term of order o* corresponding to layer *ℓ* (see Supplementary Note 1 for higher-order Taylor approximation and Supplementary Note 2 for index notation of alignment terms).

Alignment terms can provide alignment owing to their structure (Fig. 1). For simplicity, consider linear case of the alignment terms (ignoring element-wise matrix multiplications in the equation 4) where they reduce to

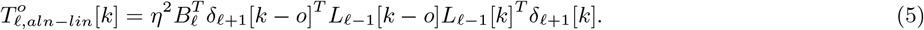

In this case, occurrence of alignment depends on the transformation matrix

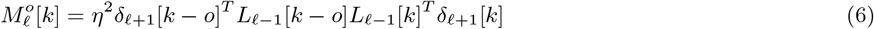

which is applied to 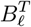 and if 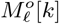 partially preserves the direction of 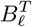 after matrix multiplication, 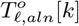 partially aligns with 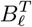. In general, 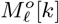 can be decomposed into its symmetric and skew-symmetric parts 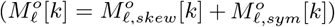. Any skew-symmetric transformation matrix totally deviates the direction and 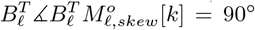 (see Supplementary Note 3). Hence, the occurrence of alignment depends on 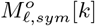.

Under certain conditions, some simplifying assumptions can be considered for the analysis of alignment terms. For example, initializing elements of 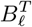 independently from *𝒩* (0, *σ*^2^) is a common practice in ANNs. If elements of 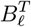 are independent random variables with a mean of zero and a variance of *σ*^2^, and if 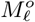 is independent of the backward weights (validity of this assumption will be discussed in a practical nonlinear ANN below), the alignment is expected if the mean of eigenvalues of 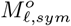 (or equivalently the mean of eigenvalues of 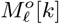) is positive. To see this, we can examine the Frobenius inner product of 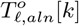 and 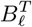, which is positive if and only if 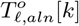 is partially aligned with 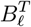, and evaluate its expected value as follows

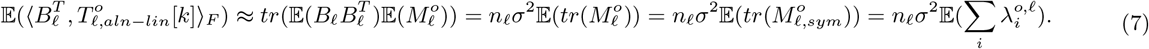

where 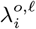 denotes the eigenvalues of 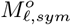. Equivalently, alignment is expected if 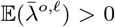 where 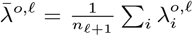. For example, in an extreme case, if 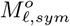 is positive semidefinite and transform an arbitrary vector, it scales each of the components of the vector with a nonnegative scalar which is the corresponding eigenvalue. In other words, it keeps each component in its previous direction and does not flip it by 180°, which is desirable for alignment. However, 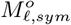 being semidefinite is not necessary and given the above assumptions, alignment is expected as long as 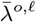 is positive. There can also be more complex conditions where 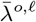 is positive, but rows of 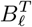 lie near some eigenvectors whose corresponding eigenvalues are negative. In this case, alignment does not occur despite having a positive 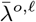. However, given a random and independent *B*_*ℓ*_ (absence of any strong dependence between *B*_*ℓ*_ and 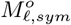) such a condition is unlikely.

The structure of alignment terms provides a robust basis for occurrence of alignment under a variety of conditions depending on the properties of neural activity. In particular, autocorrelation of error (*δ*_*ℓ* +1_) and output signals (*L*_*ℓ* −1_) of neurons and also cross-correlation between them play an important role in WA. Considering the above simplifying assumptions, the role of autocorrelation and cross-correlation can be seen by breaking 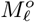 into its constituent terms as follows

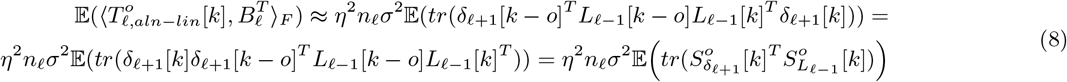

where we define 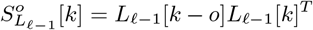 as *output similarity matrix* of layer *ℓ* − 1 and 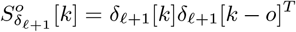 as *error similarity matrix* of layer *ℓ* + 1. These two matrices represent the autocorrelation of output and error signals of neurons across lags, and their multiplication in the equation 8 represent the cross-correlation between them.

To verify this conclusion, we performed a simulation using an open-loop 2-layer ANN with ReLU nonlinearity where we manually set the output signals of input neurons (*L*_0_) and error signals of output neurons (*δ*_2_) and controlled their autocorrelation and cross-correlation (Fig. 2A). For example, in an extreme hypothetical condition where we initially drew elements of *L*_0_ and *δ*_2_ independently from 𝒩 (0, 1) and left them constant across iterations, the transformation matrix 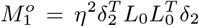 is a symmetric positive semidefinite matrix (Fig. 2B first row), and alignment happens as predicted (Fig. 2C blue trace). In this case, expected values of both 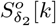 and 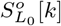 are scalar multiples of the identity matrix.

**Figure 2.**
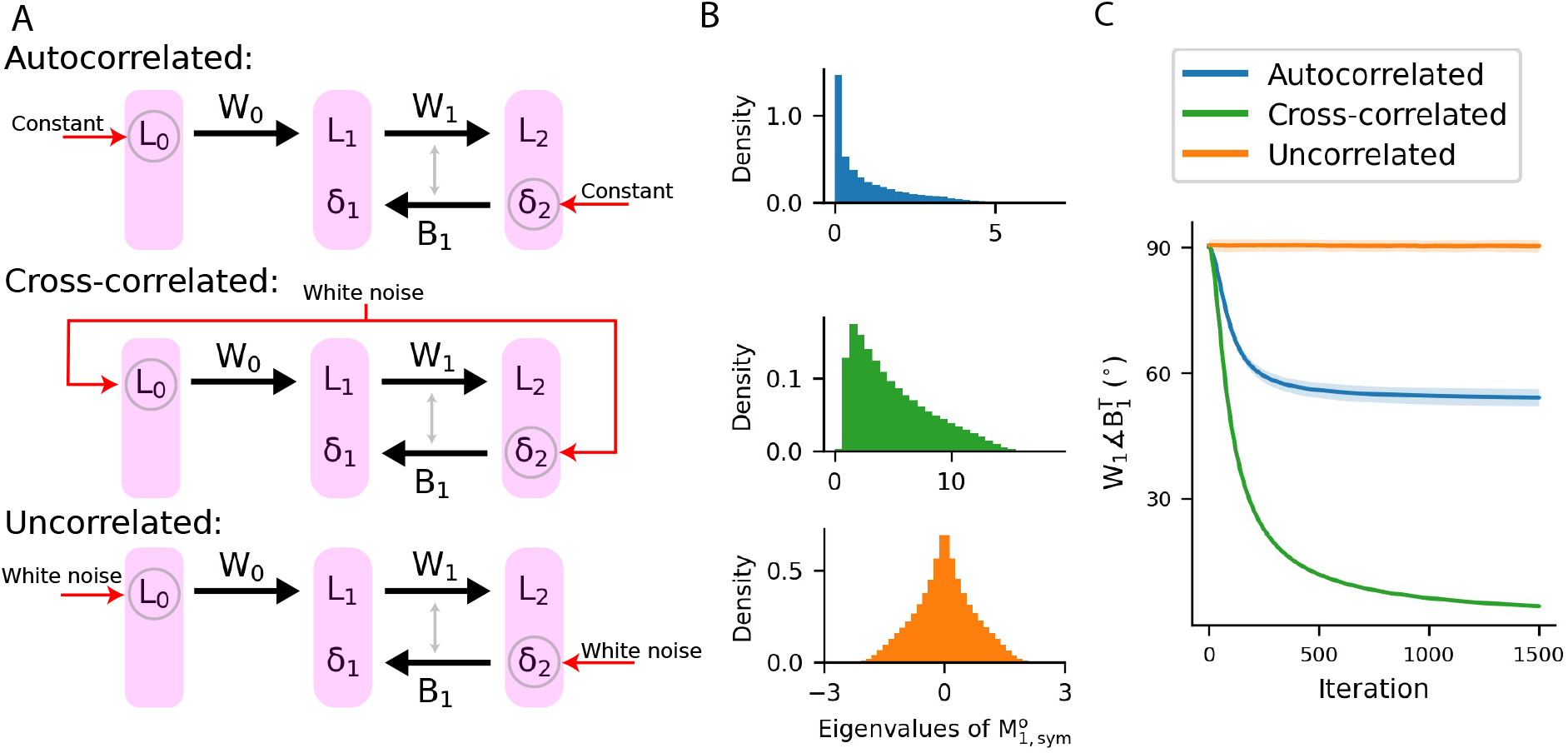
Weight alignment can occur in the absence of meaningful input and error depending on the statistical properties of neural activity. **(A)** In an open-loop nonlinear 2-layer ANN, hypothetical output signals are imposed on the neurons in the input layer (*L*_0_) and error signals of neurons in the output layer (*δ*_2_). In the top row, elements of *δ*_2_ and *L*_0_ are independently generated from 𝒩 (0, 1) at the beginning and left constant across all iterations. In this case, inputs and errors are autocorrelated. In the middle row, elements of *δ*_2_ = *L*_0_ are independently generated from *𝒩* (0, 1) at each iteration. In this case, elements of *δ*_2_ and *L*_0_ are not autocorrelated, but they are cross-correlated. In the bottom row, elements of *δ*_2_ and *L*_0_ are independently generated from *𝒩* (0, 1) at each iteration. In this case, input and error signals are neither autocorrelated nor cross-correlated. **(B)** Histograms of eigenvalues of random samples of 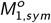 corresponding to the conditions of panel A. In the top and middle row scenarios, the mean of the eigenvalues are positive, but, in the bottom row, the mean of them is zero. **(C)** The angle between forward and backward weights 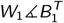 in the scenarios of panel A. Each trace is the average over 10 runs and shaded areas are one s.d. around the mean.

In another case, we re-initialized elements of *L*_0_ independently from *𝒩* (0, 1) at each iteration and let *δ*_2_[*k*] = *L*_0_[*k*]. In this case, output signals of input neurons and error signals of output neurons are white noise and are not autocorrelated, but they are fully cross-correlated. Here, although the expected values of elements of 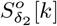 and 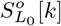 at any given lag *o ≠* 0 are zero, they are positively cross-correlated, and alignment happens (Fig. 2C, green trace). In contrast, if all elements of *L*_0_ and *δ*_2_ are independently re-initialized from *𝒩* (0, 1) at each iteration, error signals of output neurons and output signals of input neurons are neither autocorrelated nor cross-correlated and alignment does not happen (Fig. 1B, orange trace). Notably, the occurrence of alignment in these scenarios can be predicted from the distribution of the eigenvalues of 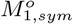 as in the last scenario with 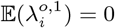 alignment does not happen (Fig. 2B).

Note that despite the simplifying assumption of linearity for the analysis of alignment terms, in these three scenarios, the network was nonlinear. Nonlinearity appears as two element-wise matrix multiplications in the structure of alignment terms (equation 4). However, using an increasing activation function, such as ReLU, causes the elements of the matrices resulted from *f*′ (·) to be nonnegative, which alleviate the impact of nonlinearity. Moreover, given the actual behavior of the nonlinear network, there is no game-changing dependence between matrices resulted from *f* ^*′*^ (·) and other signals of the network (*L*_0_ and *δ*_2_) such that invalidates the conclusion of the linear analysis. Under these circumstances, we can regard nonlinearity as a distortion that impacts the amount of alignment but does not determine its occurrence (see Supplementary Note 2), and for simplicity, we can ignore nonlinearity and refer to the linear case for the analysis of the nonlinear alignment terms (will be demonstrated in a practical nonlinear ANN below).

In DFA (Nøkland, 2016), instead of backpropagation of error signals step by step from each layer to its previous layer, error signals are backpropagated directly from the output layer to each hidden layer through direct fixed random weights, for instance, 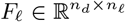. With this method, it has been reported that the product of forward weights (*W*_*ℓ*_ *W*_*ℓ* +1_ *… W*_*d*−1_) aligns with 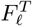 (Crafton et al., 2019). In this work, we focus on FA, but it can be shown that a similar technique with Taylor series expansion can be used to explain DFA (see Supplementary Note 4).

### 2.2 Investigating BP-TAW in training a 5-layer ANN on MNIST

To have a concrete example, in the subsequent sections, we examine the dynamics of alignment terms to investigate BP-TAW in the learning process of a 5-layer nonlinear fully connected ANN (Fig. 3A) for handwritten digits classification on MNIST dataset. For nonlinearity, we chose *f* (·) = *tanh*(*ReLU* (*·*)), which roughly resembles frequency-current curve of biological neurons. Moreover, since this is a classification task with desired output of the network coded to be between zero and one, for the reasons of stability and convergence, it is convenient for the activation function of the output layer to be confined between zero and one. We matched the number of neurons and also activation functions of all hidden and output layers so that the amount of alignment can be comparable between different layers. We chose the number of neurons in each hidden and output layer to be 50 and since there were 10 classes, to match the length of the coding of desired output of the network with the number of neurons in the last layer, we coded the labels of classes with mutually exclusive 5-hot coding (see Methods).

**Figure 3.**
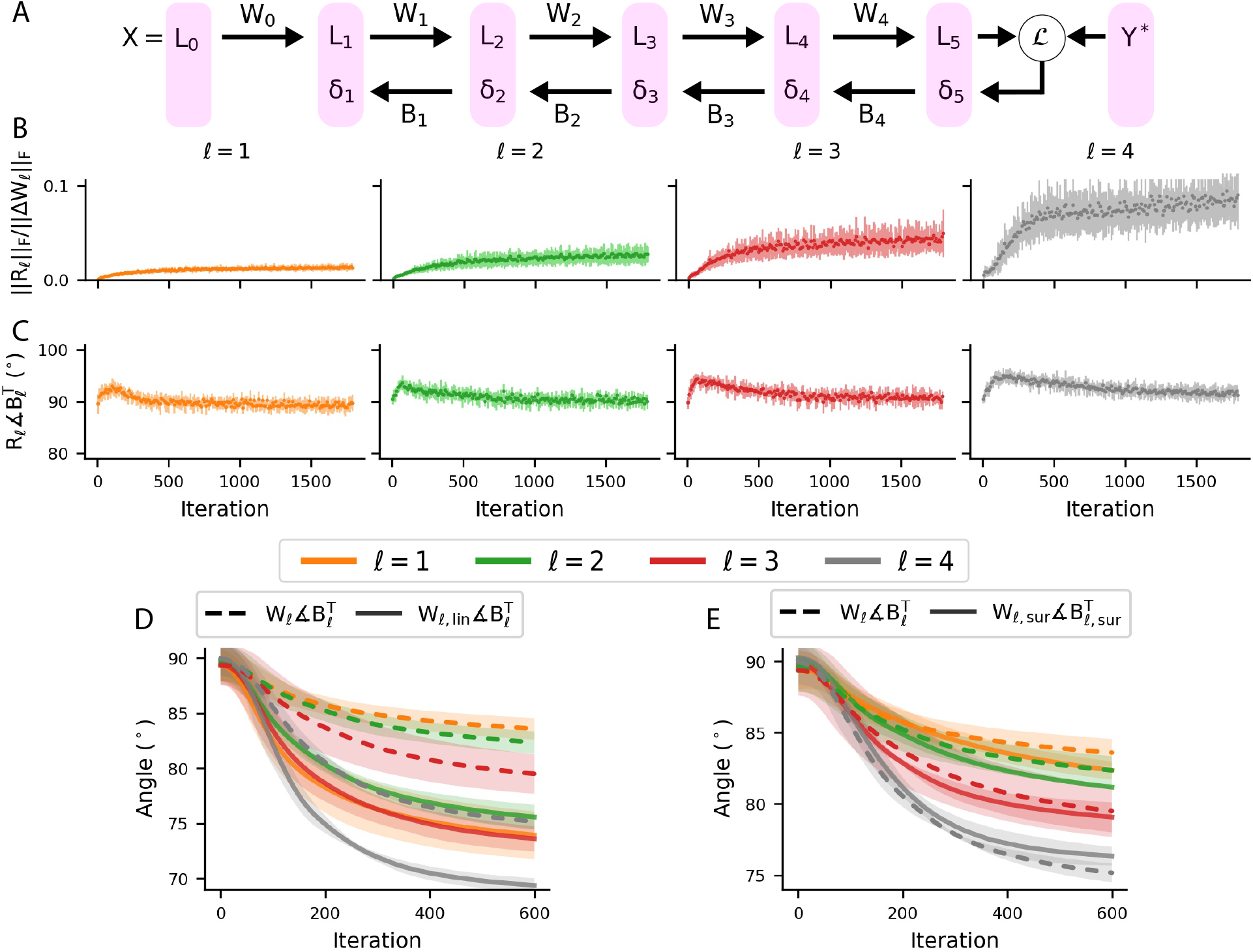
Effect of approximations and simplifying assumptions on alignment in the training of a nonlinear ANN on MNIST. **(A)** Layout of the network under investigation. **(B**,**C)** To extract the alignment terms, first order Taylor approximation is used (equation 4). In this example network, the remainder terms (*R*_*ℓ*_) constitute a small proportion of weight update directions. Moreover, they remain roughly orthogonal to the backward weights. Therefore, the remainder terms are not responsible for the occurrence of alignment and the first-order Taylor approximation provides a good approximation for the analysis of WA. **(D)** In the learning process of the network where its actual weight matrices (*W*_*ℓ*_) are updated by the update rule of BP-TAW, *W*_*ℓ, lin*_ is updated by a different version of update rule of BP-TAW (equation 10) where for calculating the alignment terms, nonlinearity is ignored (equation 5 is used instead of 4), but other matrices involved are the same as the actual network. Comparing two cases shows that nonlinearity is a distortion on the alignment terms which reduces the amount of alignment but does not prevent it from occurring. Therefore, for simplification, nonlinearity can be ignored in the analysis of WA in this example network. **(E)** In the learning process of the network where its actual weight matrices (*W*_*ℓ*_) are updated by the update rule of BP-TAW, *W*_*ℓ, sur*_ is updated by a different version of update rule of BP-TAW (equation 11) where for calculating alignment terms actual backward weights of the network are replaced with surrogate backward weights (*B*_*ℓ, sur*_), which are randomly generated, while other matrices involved are the same as the actual ones. The alignment between 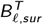 and *W*_*ℓ, sur*_ shows that this replacement slightly affects the amount of alignment but does not prevent the alignment from occurring. Therefore, in this example network, dependence between *B* and 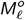 plays a minor role in WA. (B-E) Each dot or trace is the average over 30 runs and error bars and shaded areas are one s.d. around the mean.

To reduce the computational cost, we resized all handwritten digits (data points of MNIST) to images with 15 *×* 15 pixels which were then transformed into a vector. Hence, the number of input neurons was 225. We also normalized input data points (output signals of input neurons) to lie between 0 and 1 (dividing the original MNIST data points by 255). We chose the batch size to be 1000, which means there were 60 mini-batches given the total number of 60000 training data points (each epoch of training consisted of 60 iterations). We also examined the effect of data shuffling (rearrangement of data points among all mini-batches at the beginning of each epoch) and show that data shuffling has no considerable effect on total WA, while affects individual alignment terms.

At the beginning of each run, we randomly initialized elements of forward and backward weights and bias vectors independently from *𝒩* (0, 0.1). The loss function that we used was 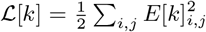, where *E*[*k*]_*i,j*_ is the element in the *i*^*th*^ row and the *j*^*th*^ column of *E*[*k*] = *Y* ^*^[*k*] − *Y* [*k*].

#### 2.2.1 Effect of approximations and simplifying assumptions on alignment

In this section, we will evaluate the effects of approximations and assumptions used in derivation and analysis of alignment terms in the example network under investigation.

##### 2.2.1.1 First-order Taylor approximation

In derivation of alignment terms for nonlinear networks (equation 3), we used first-order Taylor approximation. Namely, each order of alignment terms 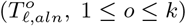 is the first-degree Taylor polynomial of successive Taylor series and we have neglected higher-degree Taylor polynomials (remainder term), which induces some error in the analysis of WA. Moreover, in practice, the activation function used may have some nonanalytic points in its domain (for example, ReLU at zero), inducing additional error in the analysis of WA. Therefore, investigating the approximation error is an important step in verification of usefulness of the alignment terms.

We define the difference between the true Δ*W*_*ℓ, F A*_ and its approximation as

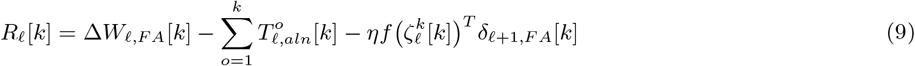

which we refer to as the remainder term of layer *ℓ*. The first-order Taylor approximation can be acceptable if *R*_*ℓ F*_ is small relative to ‖Δ*W*_*ℓ, F A*_ ‖_*F*_. It would be also interesting to see whether *R*_*ℓ*_ also aligns with 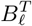. Analysis of this specific ANN shows that across iterations there is a modest increase in ‖*R*_*ℓ*_ ‖ _*F*_ relative to ‖Δ*W*_*ℓ, F A*_ ‖_*F*_, which is more pronounced in the final layers, but it remains small (*<* 10%) relative to ‖Δ*W*_*ℓ, F A*_‖_*F*_ (Fig. 3B). Angle analysis reveals that *R*_*ℓ*_ is not aligned with 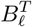 and even slightly opposes the alignment in the initial phase of the learning process (Fig. 3C), that is, remainder terms are not responsible for the occurrence of alignment and even slightly interfere with it. Hence, alignment terms are able to sufficiently explain the degree and dynamics of alignment in this network.

In addition to alignment terms, another term that appears in the Taylor series expansion is 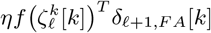 (equation 4), which is the zero-degree Taylor polynomial of the last Taylor series expansion used for extracting the alignment term of order *o* = *k*. This term does not have the beneficial structure of the alignment terms and does not show a considerable amount of alignment in this network (Fig. S1).

##### 2.2.1.2 Ignoring nonlinearity

To investigate the effect of nonlinearity on the alignment terms, in the learning process of the network by BP-TAW, in addition to the actual forward weights (*W*_*ℓ*_), we updated a different version of the forward weights (*W*_*ℓ, lin*_) with a different version of update direction as follows

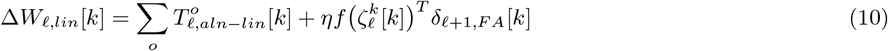

where we ignored nonlinearity in the calculation of the alignment terms (using equation 5 instead of 4), while the error and output signals of the actual intact nonlinear network were still used for the calculation of the linear alignment terms (*W*_*ℓ, lin*_ was not used for the calculation of neural activity). In order to examine the impact of nonlinearity only on the alignment terms, we preserved the nonlinearity in the zero-degree Taylor polynomial (the rightmost term in the equation 10).

Comparing the alignment of the actual forward weights with the alignment of Δ*W*_*ℓ, lin*_ shows that regardless of the amount of alignment, the overall dynamics of linear and alignment terms are similar (Fig. 3D). In other words, nonlinearity is a distortion on the alignment terms that reduces the amount of alignment but does not prevent alignment from occurring in this network.

##### 2.2.1.3 Independence of the transformation matrix 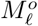 and backward weights *B*_*ℓ*_

After the start of the learning process, both the output and error signals of neurons at each layer are a function of the backward weight matrices. However, as explained in the derivation of equation 7, if the statistical dependence between the transformation matrix 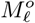 and backward weights is weak, we can mainly attribute the occurrence of WA to the properties of the transformation matrix 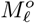.

If the alignment is driven by the properties of 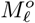 rather than the relationship between 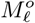 and 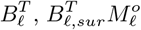 still aligns with 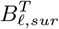, where the actual backward weights used for the training (*B*_*ℓ*_) are replaced with surrogate backward weights (*B*_*ℓ, sur*_) which are randomly generated from the same distribution of the actual backward weights. Surrogate backward weights have no contribution to the training of the network, and consequently they are independent of 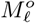.

To examine this, in the learning process of the network by BP-TAW, in addition to the actual forward weights (*W*_*ℓ*_), we updated a surrogate version of the forward weights (*W*_*ℓ, sur*_) with a surrogate version of update directions according to

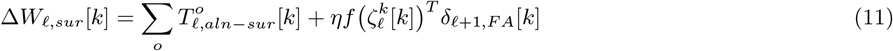

where the surrogate nonlinear alignment terms 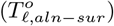 were calculated according to the equation 4, except that the actual backward weights were replaced with surrogate backward weights, while other matrices involved in the surrogate alignment terms were the same as the matrices of the actual intact network (*W*_*ℓ, sur*_ was not used for the calculation of neural activity and *B*_*ℓ, sur*_ was constant in each run).

The comparison of alignment of the actual weights 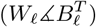 with the alignment of surrogate weights 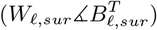 shows that a similar alignment, with a minor difference in amount, happens across layers in both situations (Fig. 3E). Thus, we can assume that alignment is largely driven by the properties of 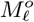 and that the statistical dependence between 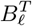 and 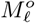 (and also matrices involved because of the effect of nonlinearity) plays a minor role in WA in this network.

#### 2.2.2 Factors affecting the dynamics of alignment terms and alignment

##### 2.2.2.1 Autocorrelation of error and output signals of neurons contributes to alignment

Without data shuffling, in the initial phase of the learning process, all orders of alignment terms showed a considerable amount of alignment, but those whose orders were a multiple of 60 were slightly more aligned than their adjacent orders (Fig. 4A, Fig. S2A). With the continuation of the learning process, the amount of alignment of the alignment terms whose orders were not a multiple of 60 decreased, while the terms whose orders were a multiple of 60 became more aligned (Fig. 4A, Fig. S2A).

**Figure 4.**
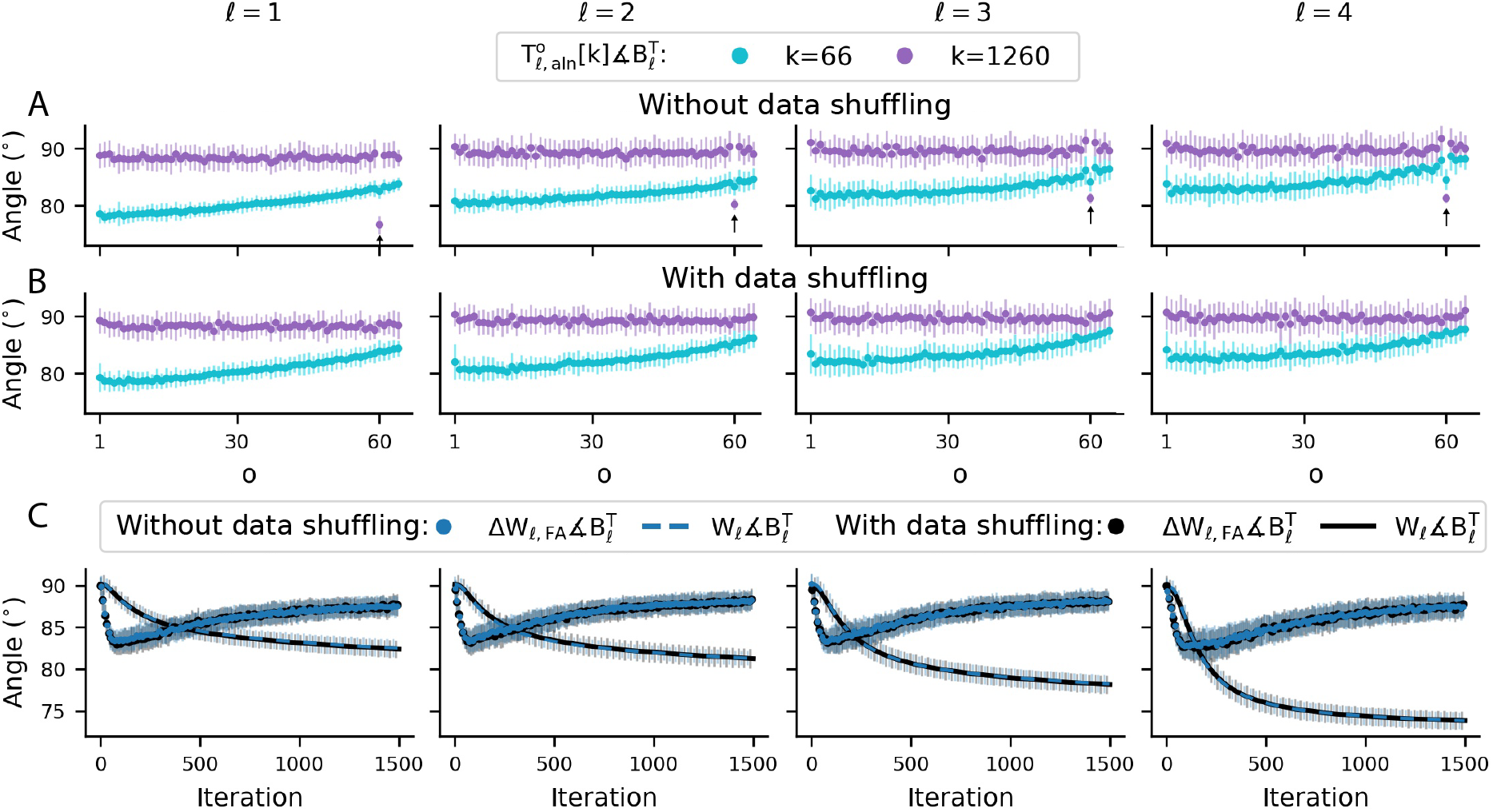
Repetition of the same data points in each epoch contributes to alignment by shaping autocorrelated neural activity. **(A)** The training dataset is divided into 60 mini-batches which are the same in all epochs (no data shuffling). As the learning process proceeds, the alignment terms whose orders are not a multiple of 60 lose their initial amount of alignment, while the one whose order is 60 becomes more aligned. This difference is due to the repetition of the same mini-batches every 60 iterations which makes neural activity autocorrelated at the lag of 60 and its multiples. **(B)** The training dataset is divided into 60 mini-batches and also is shuffled at the beginning of each epoch (the arrangement of data points across mini-batches is changed). Unlike the case without data shuffling, the alignment term of order 60 does not behave differently since the same arrangement of mini-batches is not repeated in every epoch. **(C)** Data shuffling changes the behavior of alignment terms but not the total alignment since the autocorrelated activity of neurons that without data shuffling is concentrated in the lags that are a multiple of 60, with data shuffling becomes distributed among other lags. (A-C) Each dot or trace is the average over 30 runs and error bars are one s.d. around the mean.

The arrangement of data in the mini-batches effects the behavior of alignment terms. Since the dataset is divided into 60 mini-batches, without data shuffling, the same mini-batches are repeated every 60 iterations. As a result, neural activity is autocorrelated at lags that are a multiple of 60, causing the alignment terms whose orders are a multiple of 60 to behave differently. Referring back to the equation 5, an appropriate condition for occurrence of alignment which makes 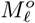 a positive semidefinite matrix is that *L*_*ℓ* −1_[*k* − *o*] and *δ*_*ℓ* +1_[*k* − *o*] are equal to *L*_*ℓ* −1_[*k*] and *δ*_*ℓ* +1_[*k*], respectively. Without data shuffling and by having a small learning rate, this condition is approximately satisfied for lags that are a multiple of 60. Moreover, as the network learns, its response to data points, that is, the output and error signals of its neurons produced by each data point, becomes more stable. Hence, in comparison to the initial phase, in the late phase of the learning process, *L*_*ℓ* −1_[*k* − 60] and *δ*_*ℓ* +1_[*k* − 60] become more similar to *L*_*ℓ* −1_[*k*] and *δ*_*ℓ* +1_[*k*], respectively, making alignment term of order 60 more aligned (Fig. 4A, Fig. S2A). With data shuffling, this considerable amount of autocorrelation does not exist in the activity of neurons at lags that are a multiple of 60, and thus alignment terms whose orders are a multiple of 60 behave similar to the other orders of alignment terms and lose their initial amount of alignment with the continuation the learning process (Fig. 4B, Fig. S2B).

However, the amount of alignment of Δ*W*_*ℓ*_ with 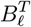 does not change with data shuffling (Fig. 4C). Assuming the update steps to be relatively small, in terms of statistical properties of neural activity, shuffling is similar to substituting the rows of error and output matrices across different lags with each other. As a result, the autocorrelated activity of neurons that without data shuffling is concentrated in the lag of *o* = 60, with data shuffling becomes distributed among the lags of *o* = 60 up to *o* = 119 since a specific data point that appears in the mini-batch of the *k*^*th*^ iteration, will inevitably be repeated in one of the mini-batches of (*k* + 60)^*th*^ to (*k* + 119)^*th*^ iteration. The alignment of *W*_*ℓ*_ with 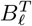 is influenced by the sum of Δ*W*_*ℓ*_ and the alignment of Δ*W*_*ℓ*_ itself is influenced by the resultant of all orders of alignment terms. Data shuffling changes the behavior of individual alignment terms but preserves the collective behavior of them and has no considerable effect on the amount of alignment (see Supplementary Note 2).

Data shuffling does not affect the overall behavior of the alignment terms whose orders are not a multiple of 60. The reason of their initial alignment and also the subsequent reduction in their amount of alignment will be discussed in the next section.

##### 2.2.2.2 Cross-correlations of error and output signals of neurons contributes to alignment

It is normally the case that data points belonging to a single category are more similar to each other than the ones belonging to different categories. This property contributes in the alignment of *W*_*ℓ*_ with 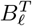 by shaping cross-correlated neural activity between *δ*_*ℓ* +1_ and *L*_*ℓ* −1_. This can be seen in the elements of similarity (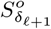 and 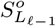). Referring back to the equation 8 and considering the aforementioned simplifying assumptions, alignment is expected if

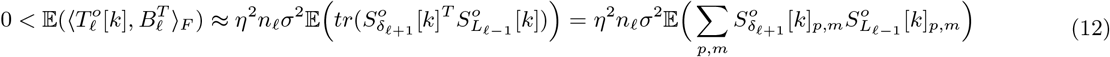

where 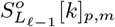 denotes the element in the *p*^*th*^ row and the *m*^*th*^ column of 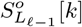. Namely, as a measure of similarity, 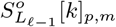 is the dot product between the output signals of neurons of the layer *R* − 1 produced by the *p*^*th*^ data point of the (*k* − *o*)^*th*^ mini-batch and their output signals produced by the *m*^*th*^ data point of the *k*^*th*^ mini-batch. 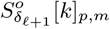 denotes a similar dot product but for the error signals of neurons in the layer *R* + 1.

In the summation of equation 12, we define 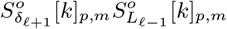 as *similarity term*. Based on the categories of the *p*^*th*^ data point of the (*k* − *o*)^*th*^ mini-batch and the *m*^*th*^ data point of the *k*^*th*^ mini-batch, the similarity terms have different behaviors. If these two mentioned data points both belong to the same category, we regard their corresponding similarity term as a *within-category similarity term*, and if they belong to two different categories, we regard their corresponding similarity term as a *between-category similarity term*.

Since the activation function used is nonnegative, output signals of neurons are always nonnegative and consequently 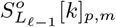 is nonnegative. In within-category similarity terms, since data points have the same true label, their error signals are mostly similar to each other and 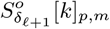 has a positive mean (Fig. 5A, Fig. S3A), which is constructive for alignment (referring to equation 12 and given that 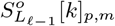 is nonnegative). In contrast, in between-category similarity terms, since data points have different true labels, their error signals are dissimilar to each other and 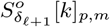 has a negative mean (Fig. 5A,Fig. S3A), which is destructive for alignment (referring to equation 12 and given that 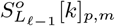 is nonnegative).

**Figure 5.**
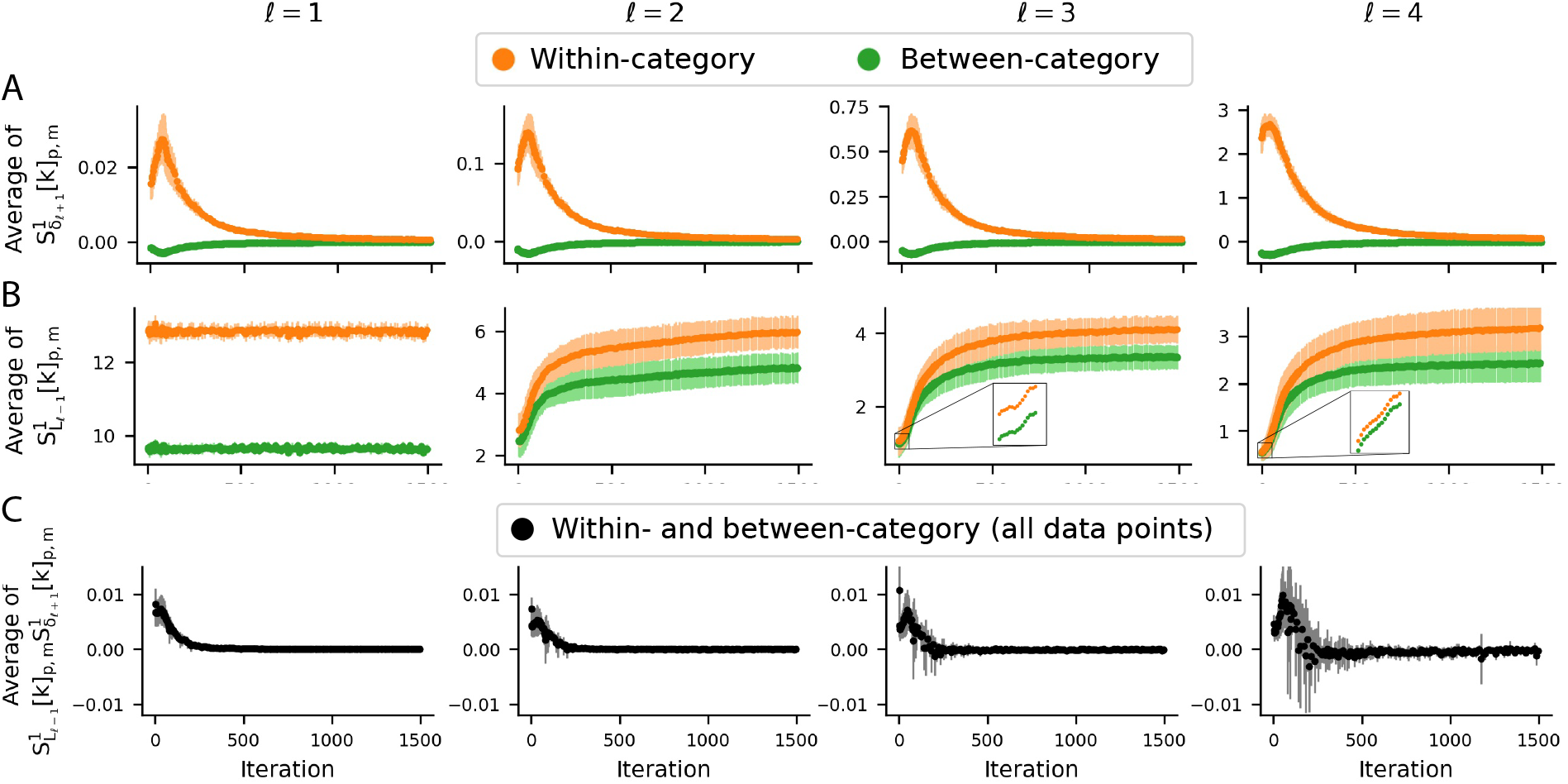
Relative similarity of data points belonging to a single category compared to data points belonging to different categories contributes to alignment by shaping cross-correlated neural activity. **(A)** As a measure of similarity, 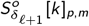 is the dot product between the error signals of the layer *ℓ* + 1 produced by the *p*^*th*^ data point of the (*k* −*o*)^*th*^ mini-batch and the *m*^*th*^ data point of the *k*^*th*^ mini-batch. Within-category corresponds to the condition that both of these two data points belong to a single category and between-category corresponds to the condition that they belong to different categories. **(B)** The same as A but for output signals of neurons. At each iteration, on average, within-category data points and also their representations across layers are more similar to each other than data points belonging to different categories. **(C)** In the early phase of the learning process, this cross-correlation between 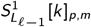 and 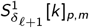 contributes to the positivity of the average of 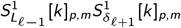 and makes the alignment to be expected (equation 12). As the learning process proceeds, the network learns to classify, and it makes the majority of error signals vanish, especially in response to the data points that are very similar to the other data points of their own category (see Fig. S3), causing cross-correlation between 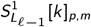 and 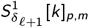 to become weaker and the average of 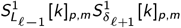 to decrease. (A-C) The averages are calculated over *p* and *m*. Each dot is the average over 10 runs and error bars are one s.d. around the mean.

However, there is an advantageous cross-correlation between 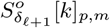 and 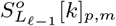 which is that in within-category similarity terms 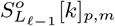 has a higher mean compared to between-category similarity terms (Fig. 5B, Fig. S3B), strengthening the constructive effect of 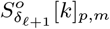 in within-category similarity terms compared to between-category ones. This cross-correlation originates from the mentioned property of the dataset which directly affects the input layer of the network and makes 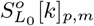 of within-category similarity terms to have a relatively high mean compared to that of between-category similarity terms (Fig. 5B, Fig. S3B). In the initial phase, although the network does not discriminate between categories, this feature is still preserved in the similarity terms of subsequent layers (Fig. 5B, Fig. S3B).

Note that in the summation of the equation 12, the number of within-category similarity terms is less than the number of between-category ones. For example, if there are 10 different categories and an equal number of data points in each of them, 10% of similarity terms are within-category and 90% of them are between-category. Nevertheless, in the initial phase of the learning process, the cross-correlation between 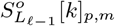and 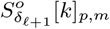 is strong enough to overcome the number and destructive effect of between-category similarity terms, and on average, 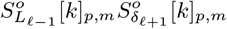 is positive (Fig. 5C).

After the initial phase, this cross-correlation becomes weaker, causing the alignment terms to lose their initial amount of alignment, except for the alignment terms whose orders are a multiple of 60 when there is no data shuffling (Fig. 4). The reason for this is that as the network learns to discriminate between different categories, the responses of neurons in the face of the majority of the data points become saturated, and error signals of them in response to them vanish (Fig. S3C), while their error signals in response to some other data points remain large (Fig. S3C). The within-category similarity of data points whose corresponding error signals remain large (and consequently their corresponding similarity terms are dominant) is less than the data points whose corresponding error signals vanish (Fig. S3D,E). This weakens the cross-correlation between error and output signals of neurons, leading to the mentioned reduction in alignment.

To verify the role of within-category similarity in deriving the alignment, we ran a simulation where we randomly selected different percentages of the data points and shuffled their true labels once at the beginning of the learning process. Such shuffling is expected to disrupt the cross-correlation between 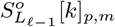 and 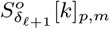 and to adversely effect the degree of WA (Fig. 6A). As expected, for high percentages of data points with randomly assigned labels, the total WA was degraded across all layers (Fig. 6A). Shuffling the labels also made it more difficult for the network to learn the classification task and increased the error rate (the test dataset was intact).

**Figure 6.**
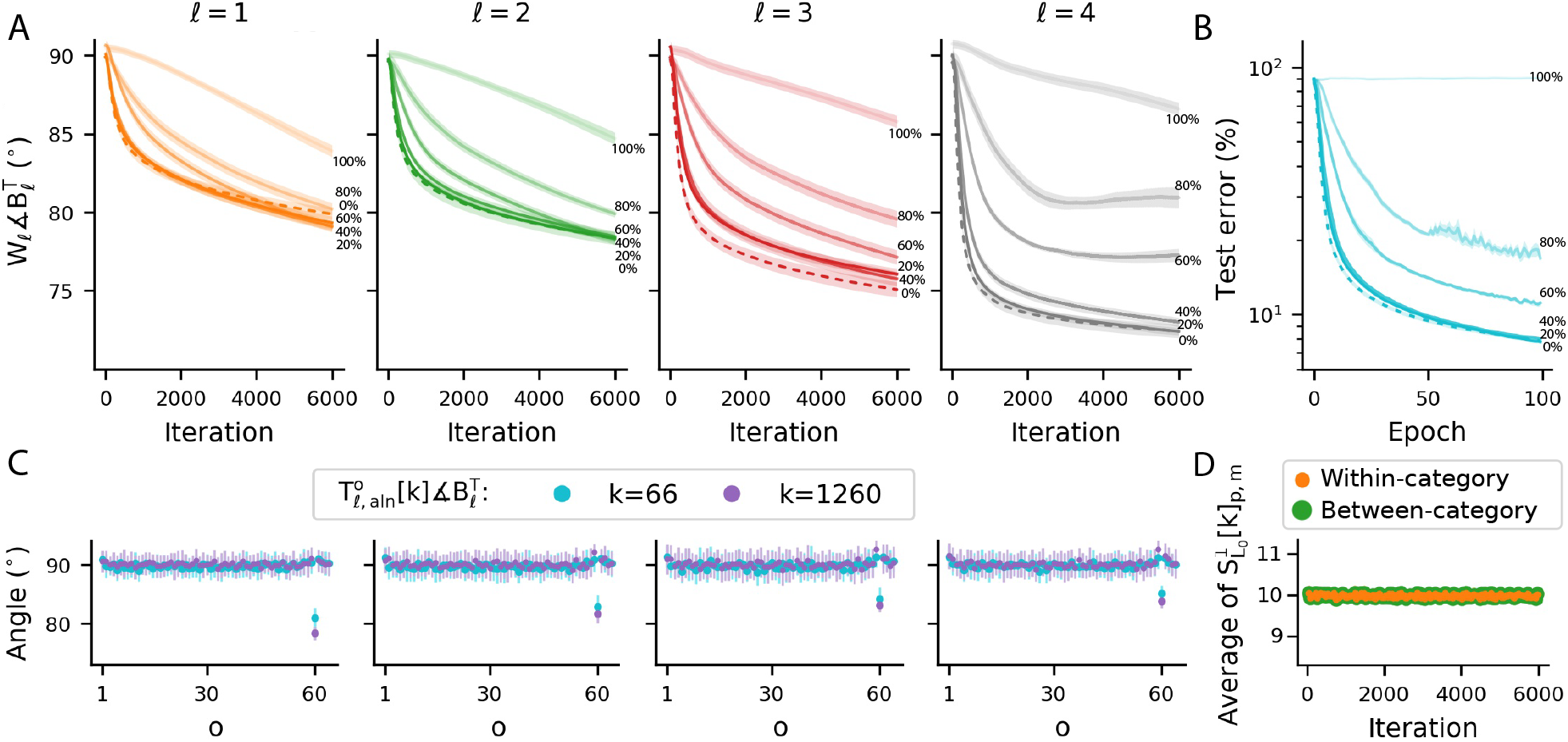
Within-category similarity of data points enhances weight alignment, but certain amount of weight alignment can happen even using data with random labels. **(A)** WA across layers of the example network trained on the MNIST dataset where labels of different percentages of data points are shuffled (in 20% label shuffling, 20% of the data points are randomly selected and then available label options are randomly assigned to them with equal probability). Alignment occurs even when 100% of the data labels are shuffled. **(B)** Classification error of the same ANN across different percentages of label shuffling. As expected, error increases for higher percentages of label shuffling and remains at the chance level (90%) if 100% of data labels are shuffled. **(C)** The behavior of alignment terms for two sample iterations *k* = 66 and *k* = 1260 when labels of 100% of the data points are shuffled once at the beginning of the learning process, and after that no data shuffling is performed. Shuffling all labels spoils the cross-correlation between error and output signals of neurons and consequently spoils alignment of all orders of alignment terms, except for the alignment term of order 60 because of the repetition of the same mini-batches every 60 iterations (compare this panel with Fig. 4A). **(D)** The average of 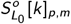 for within- and between-category data points in the case of shuffling 100% of data labels. Shuffling all labels makes the data points belonging to different categories as similar as the data points belonging to a single category and consequently spoils the cross-correlation between error and output signals of neurons (compare this panel with the leftmost column of Fig. 5B). (A-D) Each dot or trace is the average over 10 runs and error bars and shaded areas are one s.d. around the mean.

In the case where 100% of data were randomly labeled (samples of a given digit were equally likely to have one of the labels from 0 to 9) error rate reached 90%, representing the chance level (Fig. 6B). However, there was some residual and slow WA even when all labels were shuffled (Fig. 6A). The reason for the impact of label shuffling becomes clear by looking at the behavior of alignment terms across lags. In the case of shuffling 100% of the data labels and having no data shuffling at the beginning of epochs, all orders of alignment terms, except for the orders that are a multiple of 60, did not align considerably (Fig. 6C). Alignment terms whose orders were a multiple of 60 remained aligned because of the repetition of identical mini-batches every 60 iterations. In this case, 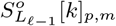 of both within- and between-category data points had the same distribution and average (Fig. 6D), and the feature that data points belonging to a single category are more similar to each other than those belonging to different categories was spoiled.

##### 2.2.2.3 Alignment and the local minimum reached by BP-TAW can be improved by weight normalization

In the training of the network with the original formulation of BP-TAW, the Frobenius norms of the forward weight matrices continuously grew especially in the last layers (Fig. S4). Unlike what is previously thought (Lillicrap et al., 2016), WA is not a direct consequence of this growth, and it can also contribute to the weakening of alignment by saturating the outputs of neurons and vanishing of error signals. To examine this, we limited and fixed the Frobenius norm of input weights to each neuron at each iteration by applying a WN method as follows

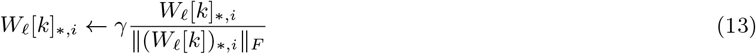

where *W*_*ℓ*_ [*k*]_*,*i*_ denotes an *n*_*ℓ*_ *×* 1 matrix that consists of the *i*^*th*^ column of *W*_*ℓ*_ [*k*] and *γ* is a positive scalar which we refer to as WN gain. To treat all weights in the same way, we also applied this WN method to backward weights once at the beginning of the learning process. Unlike the conventional WN method in ANNs (Salimans and Kingma, 2016), which is a reparameterization of BP formula, this proposed WN method is an intervention in the BP-TAW formula, which partially prevents the saturation of neurons and vanishing of error signals (Fig. S5).

Two aspects of the network that directly and simultaneously affect the amount of alignment are 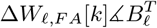 and ‖Δ*W*_𝓁,*F A*_[*k*]‖_*F*_ */*‖*W*_𝓁_ [*k*]‖_*F*_. This WN method improved both of these aspects (Fig. 7A,B) and consequently improved alignment of forward weights with backward weights (Fig. 7C) and also the alignment between update directions of BP-TAW and those of BP (Fig. 7D), making BP-TAW a better approximation of BP. In addition to the improvement of alignment, this WN method also improved test accuracy when the network was trained by BP-TAW (Fig. 7E).

**Figure 7.**
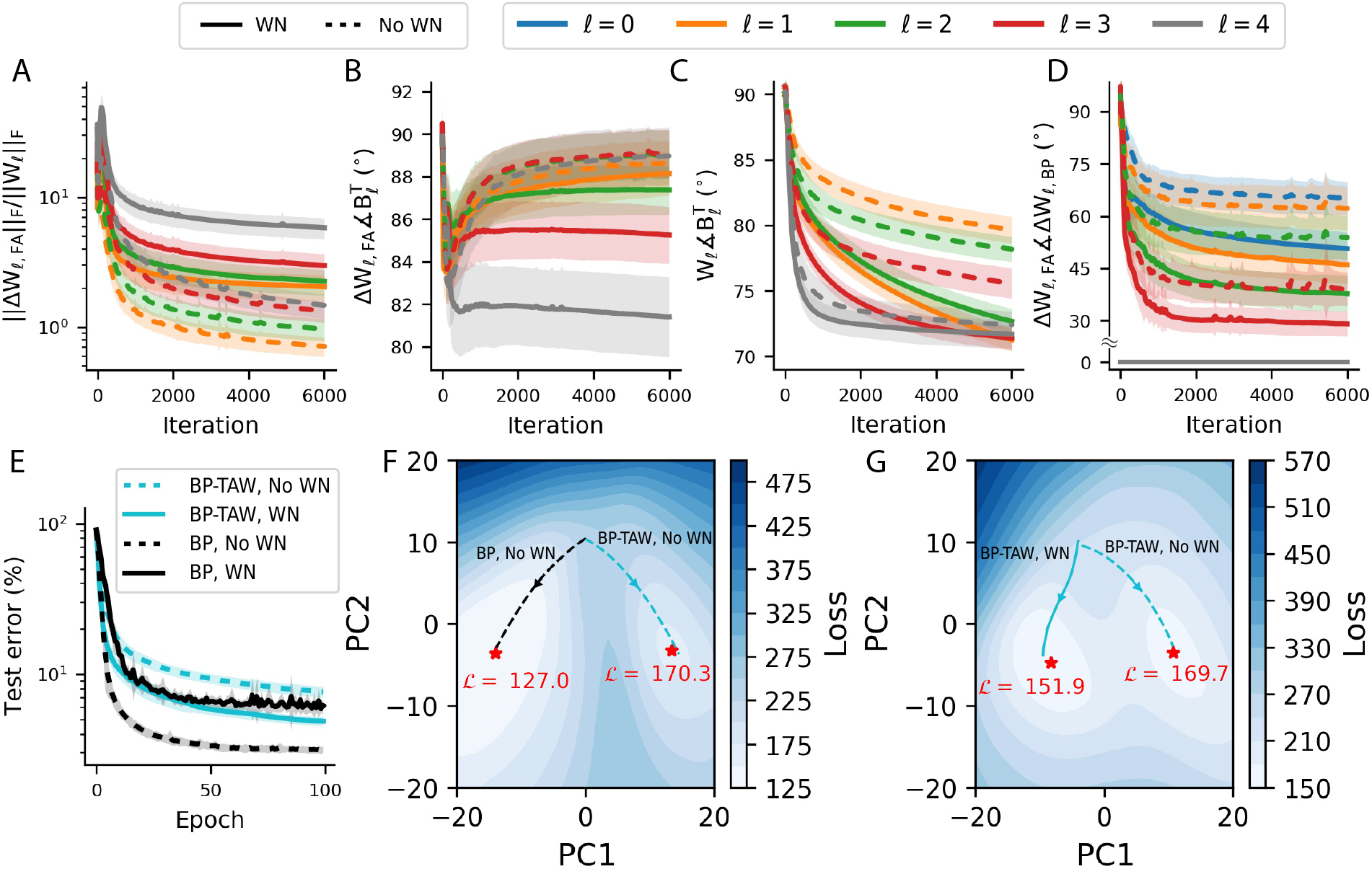
Weight normalization in BP-TAW improves alignment and classification accuracy. **(A)** WN increases the Frobenius norms of update directions of BP-TAW relative to the Frobenius norms of forward weights. **(B)** WN improves the alignment of update directions of BP-TAW with backward weights. **(C)** WN improves WA. **(D)** WN improves the alignment between update directions of BP-TAW and those of BP. In the last layer, update direction of BP-TAW is the same as BP. **(E)** WN with the gain of *γ* = 1 can improve the test accuracy of BP-TAW whereas it makes the performance of BP less robust. **(F)** Two dimensional embedding of the trajectories of all learnable parameters of the network (forward weight matrices and bias vectors) for two instances of the network that are both identically initialized with the same parameters, but one is trained by BP and the other by BP-TAW. The contour map shows the loss function of the network using the reconstructed parameters. Red asterisks are local minimums in the two dimensional space. **(G)** Similar to the panel C but for networks trained by BP-TAW, with and without WN. (A-E) Each trace is the average over 10 runs and the shaded areas are one s.d. around the mean. (A,B,D) Each trace is passed through a moving average filter with a length of 60.

The improvement of WA and the test accuracy is sensitive to *γ* and *η*. We did not optimized them and simply chose *γ* = 1 and *η* = 0.0005 based on a sensitivity analysis and grid search (Fig. S6), which showed that improvement of alignment and test accuracy of BP-TAW is robust for a fairly large range of *γ*. We also tried this WN method in training the network with BP which showed that among tested values of *γ* and *η*, the test accuracy of BP is not considerably improved by this WN method and declines in the most cases (Fig. S6).

Low dimensional embedding of all learnable parameters of the network (forward weight matrices and bias vectors) using principle component analysis showed that the initial mismatch between update directions of BP-TAW and BP drives the trajectory of the parameters of BP-TAW into a different local minimum which is less optimal than the local minimum to which the network converges with BP (Fig. 7F). WN drives the trajectory of parameters to a more optimal local minimum, resulting in a better degree of classification accuracy (Fig. 7G).

We observed that without WN, alignment between *W*_𝓁_ and 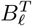 decreases in the early layers of the network. With WN, this reduction was overcome to some extent, and the first and third layers became more aligned than the last layer (Fig. 7C). However, we observed that the amount of alignment between update directions of BP-TAW and BP decreases step by step as the error is backpropagated towards the input layer and even WN could not overcome this effect (Fig. 7D). This behavior has also been reported in previous works (Moskovitz et al., 2018).

The potential increase of Δ*W*_𝓁,*F A*_∡Δ*W*_𝓁,*BP*_ (less alignment) in the earlier layers can be seen by comparing 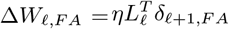 with 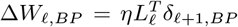. The matrix 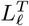 is identical in both and the factors which determine the angle between them are *δ*_𝓁+1,*BP*_ and *δ*_𝓁+1,*F A*_. For simplicity, consider a *d*-layer linear ANN. For the last layer (*ℓ* = *d*), we have *δ*_*d,F A*_ = *δ*_*d,BP*_, but for 0 *<* 𝓁 *< d*, by using the equation 2 successively, we have

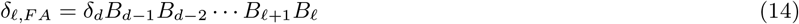

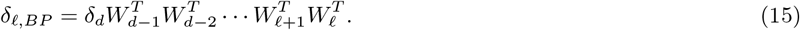

According to these two successive matrix multiplications of backward and transpose of forward weight matrices, as the error is backpropagated towards the early layers, depending on the pairs of *B*_𝓁_ and 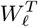, deviation of *δ*_𝓁,*BP*_ from *δ*_𝓁*F A*_ potentially tends to increase. Consequently, deviation of Δ*W*_𝓁,*F A*_ from Δ*W*_𝓁,*BP*_ potentially increases as well and it reduces the accuracy of update directions of BP-TAW compared to gradient directions computed by BP in the early layers of deep ANNs.

## 3 Discussion

Artificial neural networks and their learning paradigms have differences and similarities with biological neural networks. Specifically, BP method needs a biologically implausible matching between feedforward and feedback synaptic weights. The BP-TAW learning method (Lillicrap et al., 2016) showed that an ANN can be trained with arbitrary feedback weights that are distinct from feedforward weights. In BP-TAW, forward weights partially align with backward weights during iterations, which leads to a partial alignment between update directions of BP-TAW and BP and provides an approximation of BP.

### 3.1 Mathematical basis of alignment

In this work, we demonstrated mathematical and statistical basis of WA (Fig. 1) and showed that WA is not due to the learning or reduction of loss function; rather, it relies on the structure of alignment terms that can be extracted from the update rule of BP-TAW (equation 3 and 4), and because of this structure, alignment happens robustly under a variety of conditions depending on the neural activity. Hence, statistical properties of neural activity contribute to the occurrence of alignment and its amount. Specifically, we showed that the autocorrelation of error and output signals of neurons and the cross-correlation between them are two important features of neural activity contributing to alignment (Fig. 2).

In this work, we used alignment terms as a tool to analyze BP-TAW in a specific 5-layer nonlinear ANN trained on the MNIST dataset. We showed how the arrangement of data in mini-batches and the repetition of data points across epochs contribute to the behavior of individual alignment terms by shaping autocorrelated neural activity. Moreover, we showed that relative similarity of data points of a single category and their differences across categories, which is an intrinsic property of datasets, contributes to alignment by shaping cross-correlated neural activity.

In the analysis of BP-TAW in the specific ANN above, we considered some approximations and simplifying assumptions. We used first-order Taylor approximation to extract alignment terms, ignored the effect of nonlinearity on them, and assumed their behavior to be independent of backward weights. These approximations and assumptions facilitated the analysis and enabled us to easily identify the main factors involved in alignment in a nonlinear ANN. Importantly, we showed that these assumptions and approximations led to qualitatively valid predictions about WA (Fig. 3).

In general, many aspects of neural activity influence the behavior of alignment terms and make them act differently in different situations. For example, the architecture of network (activation function, number of neurons in layers, number of layers, loss function, normalization methods, etc.), hyperparameters (learning rate, batch size, etc.), and properties of dataset affect neural activity and consequently the behavior of alignment terms. However, we have provided a useful framework for investigating BP-TAW in the learning process of ANNs under various conditions which can be used to further our understanding of the FA and occurrence of WA in BP-TAW.

A weakness of BP-TAW as an approximation of BP in deep ANNs, which we mentioned in this work, is that the amount of alignment between update directions of BP-TAW and BP potentially tends to decline as the error is consecutively backpropagated towards the earlier layers (Fig. 7C). In other words, with BP-TAW, early layers potentially receive less accurate supervised error signals compared to the final layers. This potential decline may be overcome by unsupervised learning under certain conditions, which can be the subject of future work. Indeed, many aspects of the activity of neurons in lower areas of the visual system are demonstrated to be attainable with unsupervised learning models (Olshausen and Field, 1996; Barlow et al., 1961) and also there are suggestions of efficient network architectures where an ANN trained in an unsupervised manner is followed by a supervised classifier (Kheradpisheh et al., 2018).

### 3.2 Weight normalization can improve alignment

The correspondence of normalization methods in ANNs and biological ones has already been considered (Shen et al., 2020). In the context of BP-TAW, the utility of using normalization methods has been reported previously (Liao et al., 2016; Moskovitz et al., 2018). There are also numerous reports of normalization mechanisms in biological neural networks working to regulate the activity of neurons and limit the dynamic range of synaptic weights (Chistiakova et al., 2015; Bi and Poo, 1998). Moreover, in the biological neurons, homeostatic mechanisms, which regulate activity of neurons and prevent them from having high or low firing rates, are well documented (Murthy et al., 2001; Surmeier and Foehring, 2004; Turrigiano, 2012).

Accordingly, we examined a WN method, which worked by fixing the Frobenius norm of input weights of each neuron to some constant (*γ*). We showed that this WN method can improve alignment and the classification accuracy of the network (Fig. 7). Indeed, various forms of plasticity, such as heterosynaptic plasticity, are reported which regulate synaptic weights in a competitive manner, in which the potentiation of one synapse can result in the depression of other synapses to keep overall synaptic strengths under control, similar to our examined WN scheme (Chistiakova et al., 2015; Turrigiano, 2017).

### 3.3 Approximation of BP in biological networks and its further biological implausibilities

WA in the sense of reducing the angle between *W*_𝓁_ and 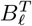 is equivalent to having positively correlated reciprocal connections between neurons in the consecutive layers. Furthermore, in a sparse network where most of the elements of *W*_𝓁_ and *B*_𝓁_ are zero, achieving greater alignment requires reciprocal connections to occur more than nonreciprocal ones. Interestingly, such a nonrandom reciprocal connectivity, is reported among pyramidal cells (Song et al., 2005; Markram et al., 1997; Holmgren et al., 2003; Sjöström et al., 2001). Although these nonrandom features are reported in local cortical circuits, if we consider synaptic plasticity as their origin (Song et al., 2005) and generalize them to the connections between neurons in different cortical layers and areas, these features can provide a favorable condition for WA. In this regard, the emergence of dominant reciprocal connections by applying spike-timing-dependent synaptic plasticity rules has been reported in a recurrent ANN of spiking neurons under rate-coded input (Clopath et al., 2010).

However, weight transport problem is only one of the biological implausibilities of BP formula which can be avoided by using BP-TAW. There are other biological implausibilities in BP and BP-TAW (Marblestone et al., 2016; Stork, 1989). For instance, firing rate as the output of each biological neuron is nonnegative, while error signals in BP and BP-TAW are signed. In addition, error signals in BP and BP-TAW are distinct from the output of artificial neurons. In BP-TAW and BP, error signals are internal attributes of neurons that are backpropagated to other neurons through feedback weights, whereas in biological networks, the attribute of neurons that is conveyed explicitly to other neurons by axons and synapses is their output spikes and it is believed that other internal attributes of them are mostly local (Stork, 1989; Song et al., 2020).

### 3.4 Conclusion

In summary, the analysis done in this study provides a useful framework for understanding WA in BP-TAW and paves the way for further research on the relationship between learning methods used in ANNs and learning mechanisms in the nervous system. While BP-TAW is capable of approximating the weight update directions proposed by BP in simple feedforward networks and WN can improve this approximation, it remains to be seen how the addition of other biological considerations such as lateral connections, sparsity, synaptic pruning and formation, and the segregation of excitatory and inhibitory neurons affect the performance of BP-TAW.

## 4 Methods

### 4.1 BP and BP-TAW learning methods

In BP, we updated bias vectors and weight matrices at each iteration as below

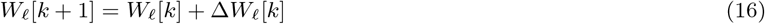

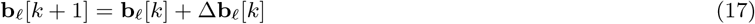

where gradient directions computed by BP for updating bias vectors and weight matrices at each iteration *k* are

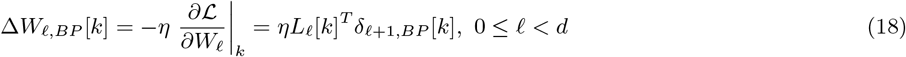

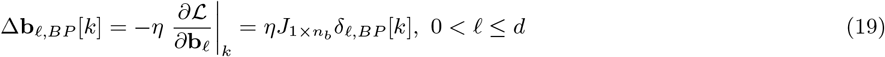

where 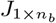 is a 1 *× n*_*b*_ all-ones matrix and error matrices of neurons are

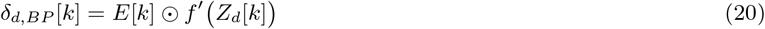

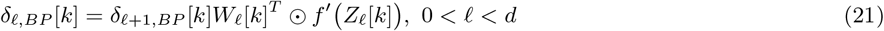

where *E*[*k*] = *Y* ^*^[*k*] − *Y* [*k*] according to the loss function 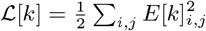 (Rumelhart et al., 1985).

In BP-TAW (Lillicrap et al., 2016), the error is backpropagated through constant random matrices different from forward weights which are denoted by 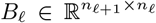, and we calculated update directions at each iteration as follows (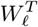 in equation 21 is replaced with *B*_𝓁_)

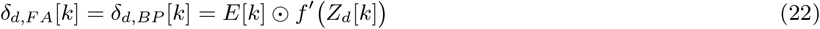

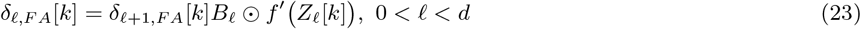

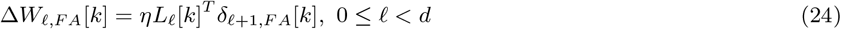

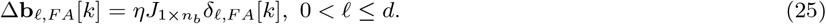

In training ANNs with BP-TAW, Δ*W*_𝓁, *BP*_ [*k*] is a direction that we only calculated at each iteration for the purpose of comparison with Δ*W*_𝓁, *F A*_[*k*] (we only used Δ*W*_𝓁, *F A*_[*k*] to update forward weight matrices).

### 4.2 Angle and cosine similarity between two matrices

We calculated the angle between two arbitrary matrices *W* and *B*, which have the same dimensions, as follows

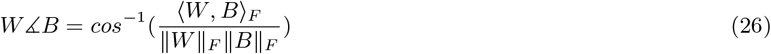

where ⟨*W, B*⟩ _*F*_ is the Frobenius inner product of *W* and *B* and ║. ║ _*F*_ is the Frobenius norm. This is identical to the angle between vectorized *W* and *B* in the euclidean space. The angle between two matrices is indeed a measure of the similarity between the normalized versions of the two matrices.

In addition to the angle, cosine similarity between two matrices can also be used as a measure of the similarity between them as follows

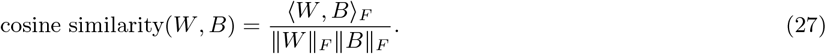

Since the denominator of the cosine similarity is always nonnegative and also assuming *W* and *B* to be nonzero, for alignment (*W ⊾ B <* 90^*°*^ or equivalently 0 *<* cosine similarity(*W, B*)) it is sufficient and necessary that

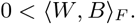

### 4.3 Network parameters, dimensions, and initialization

In Fig. 2 we chose network dimensions to be *n*_0_ = *n*_2_ = 20, *n*_1_ = 100, and *n*_*b*_ = 100, and we set *η* = 0.0004 and initialized elements of *B*_1_, *W*_0_ and *W*_1_ with i.i.d. random variables from 𝒩 (0, 1). In all experiments of training a deep ANN on the MNIST dataset by BP-TAW (with and without WN and with and without data shuffling), we set *η* = 0.0005.

### 4.4 Generating mutually exclusive n-hot coding

In training the ANN on MNIST, we used mutually exclusive 5-hot coding. Suppose the number of categories is *C* and the number of output neurons is *m* (*n · C* ≤ *m*). For generating mutually exclusive *n*-hot code vectors of size *m* for each category, we started from the first category to the last one and successively for each category *c* ∈ *{*0, 1, *· · ·, C* − 1} we initialized its code vector with zero elements and then randomly selected *n* out of *m* − *c · n* elements that were not equal to 1 in any of the *c* previously coded category vectors and set them equal to 1.

## 5 Data and Code Availability Statement

The MNIST dataset can be found at: http://yann.lecun.com/exdb/mnist/. The code for reproducing all results in this work is available under the Apache 2.0 license at https://github.com/ARahmansetayesh/FeedbackAlignmentWithWeightNormalization. We used PyTorch library only for accelerating computations on GPU (we did not use PyTorch’s automatic differentiation capability).

## 6 Author Contributions

AR, AG and FM conceived the general ideas and the research plan. AR did the derivations and simulations in discussions with and under supervision of AG. AR and AG wrote the paper under FM supervision.

## Supplementary Materials

**Figure S1.**
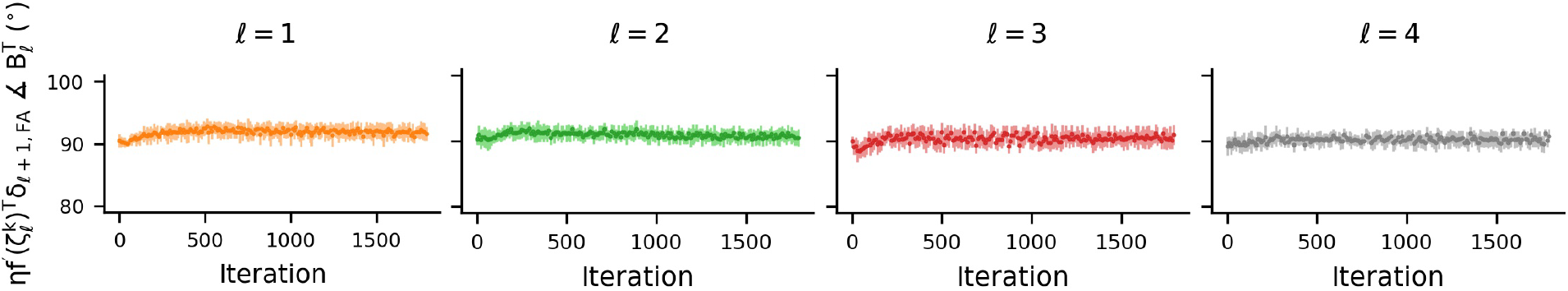
Zero-degree polynomial of Taylor approximation in the example network trained on MNIST. **(A)** In addition to the alignment terms and the remainder terms, another term that appears in Taylor series expansion is 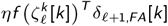 which is the zero-degree Taylor polynomial of the last Taylor series expansion applied for extracting the alignment term of order *o* = *k*. This term does not have the beneficial structure of alignment terms and does not show considerable amount of alignment. Each dot is the average over 30 runs and error bars are one s.d. around the mean.

**Figure S2.**
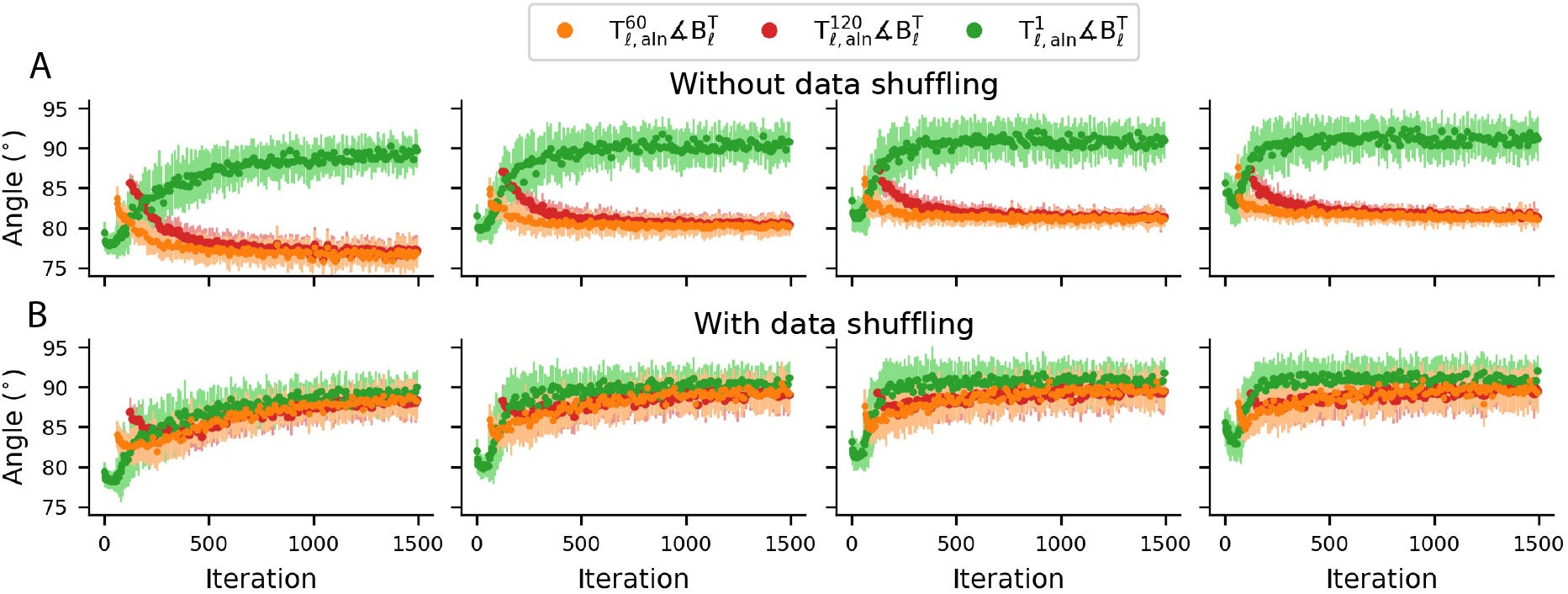
Behavior of alignment terms in the example network trained on MNIST. **(A)** The behavior of alignment terms of orders 1, 60, and 120 across iterations without data shuffling. **(B)** The same as A but with data shuffling. (A,B) Each dot or trace is the average over 30 runs and error bars are one s.d. around the mean.

**Figure S3.**
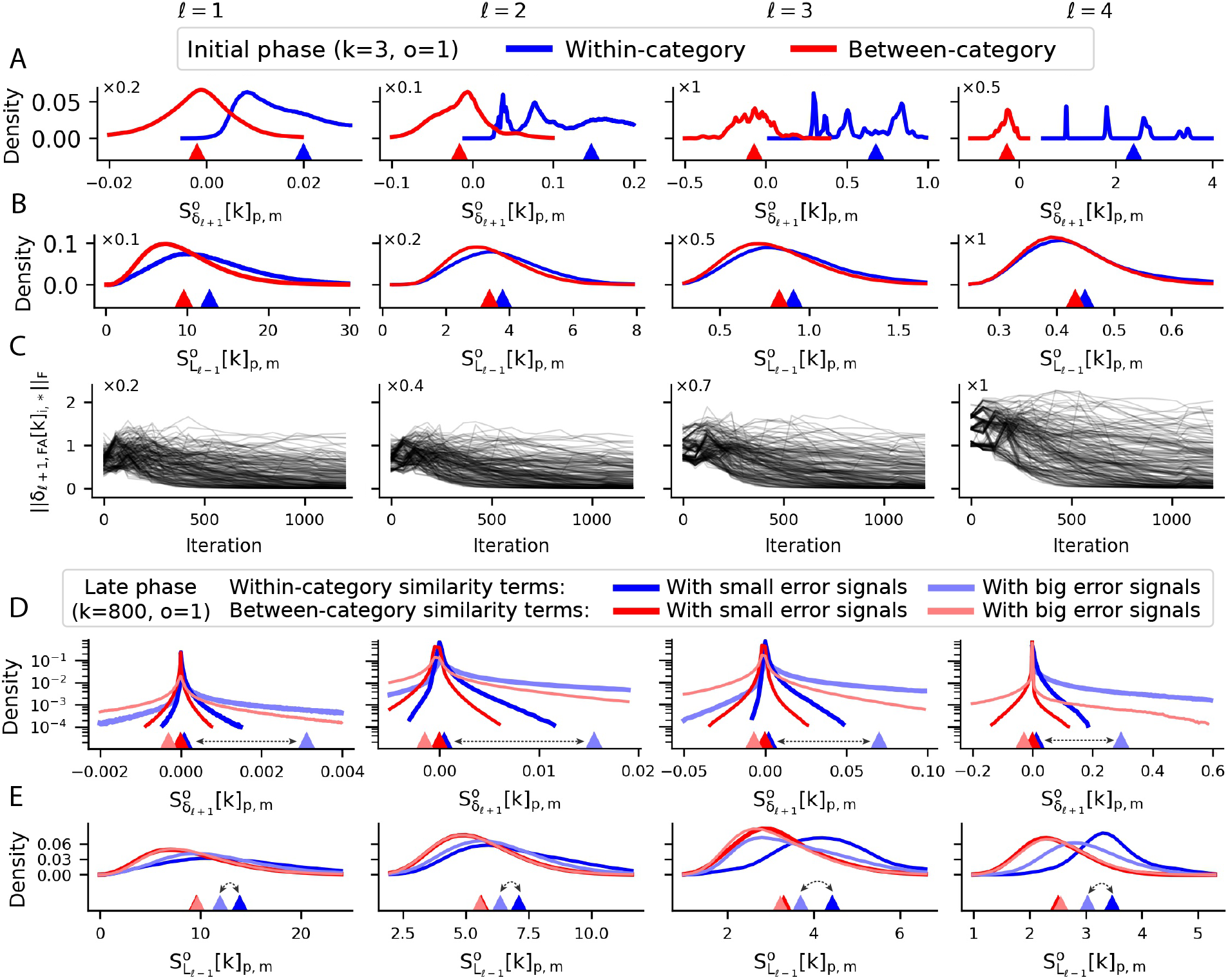
Distributions of 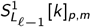 and 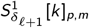 in between- and within-category similarity terms in the early and late phases of the training in the example network trained on MNIST. **(A**,**B)** Histograms of 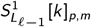 and 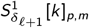 in the early phase of the network. Within-category 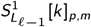 has a higher mean compared to that of between-category 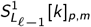 since within-category data points are more similar to each other than between-category ones. This feature shapes cross-correlated neural activity which contributes to alignment. **(C)** 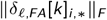 denotes the Frobenius norm of the *i*^*th*^ row of 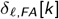 corresponding to *i*^*th*^ data point in the *k*^*th*^ mini-batch. Each trace corresponds to the Frobenius norms of error signals of a single data point of the MNIST dataset in a single run. After the initial phase, outputs of neurons in response to the majority of the data points become saturated. Consequently, the Frobenius norm of their corresponding error signals decreases to about zero and vanishes. In contrast, error signals of some data points still remain large. **(D**,**E)** Histograms of 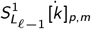 and 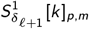 in the late phase of the network. In addition to categories, similarity terms are separated base on the magnitude of Frobenius norms of the error signals of the *p*^*th*^ data point of the (*k* − *o*)^*th*^ mini-batch and the *m*^*th*^ data point of the *k*^*th*^ mini-batch. If both of these two mentioned data points belong to a subset consisting of 60% of the total data points whose error signals at the output layer of the network have the least Frobenius norms, their similarity terms are regarded as “with small error signals”, otherwise their corresponding similarity terms are regarded as “with big error signals”. In the late phase of the network, on average, 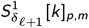 in the similarity terms with small error signals has relatively small magnitude (absolute value) compared to 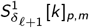 in similarity terms with big error signals (dark vs. light triangles). Thus, similarity terms with big error signals dominate the ones with small error signals in shaping the dynamics of the alignment terms in the late phase of the network. Within each category, in comparison to data points with small error signals, data points with big error signals (and also their representations across the layers of the network) are less similar to other data points of that category (dark vs. light blue triangles). This weakens the cross-correlation between 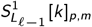 and 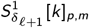 and causes 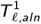 to lose its initial amount of alignment as the learning process proceeds. (A-E) Each histogram is obtained by giving the whole MNIST data points to the network. Triangles indicate the mean of each histogram. Histograms are plotted for *o* = 1, but a similar analysis holds for other orders of alignment terms.

**Figure S4.**
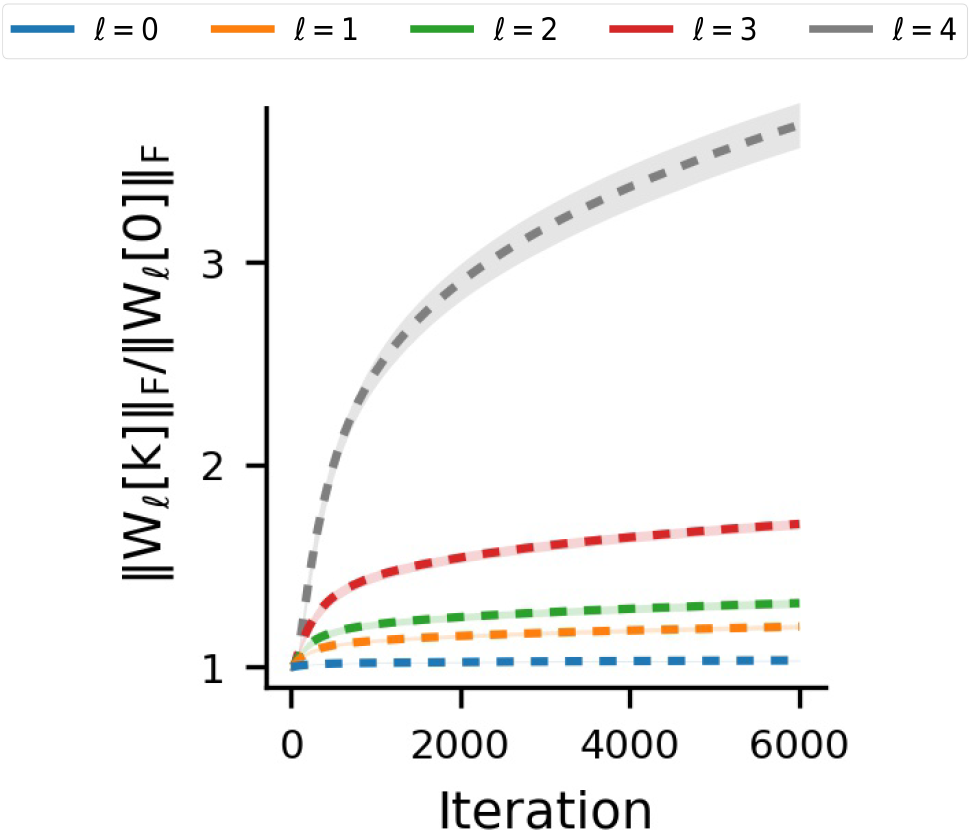
Growth of Frobenius norms of forward weight matrices in the example network trained on MNIST. Without WN, Frobenius norms of forward weight matrices continuously grow. This growth is more pronounced at the last layers.

**Figure S5.**
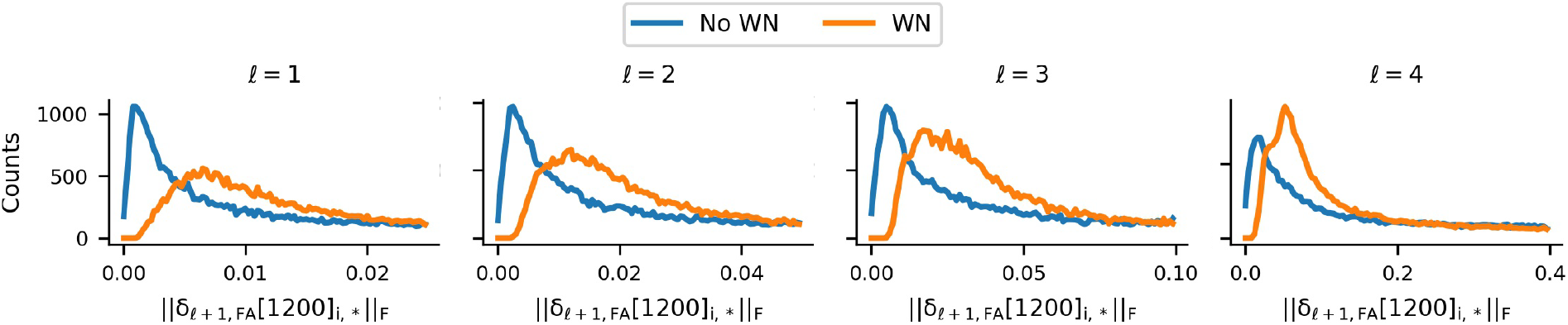
WN partially prevents saturation of neurons and vanishing of error signals in the example network trained on MNIST. Histograms of Frobenius norms of error signals of all individual data points of the dataset with and without WN across all layers. *δ*_𝓁+1,*FA*_[*k*]_*i*,*_ denotes the *i*^*th*^ row of *δ*_*𝓁* +1,*FA*_ [*k*].

**Figure S6.**
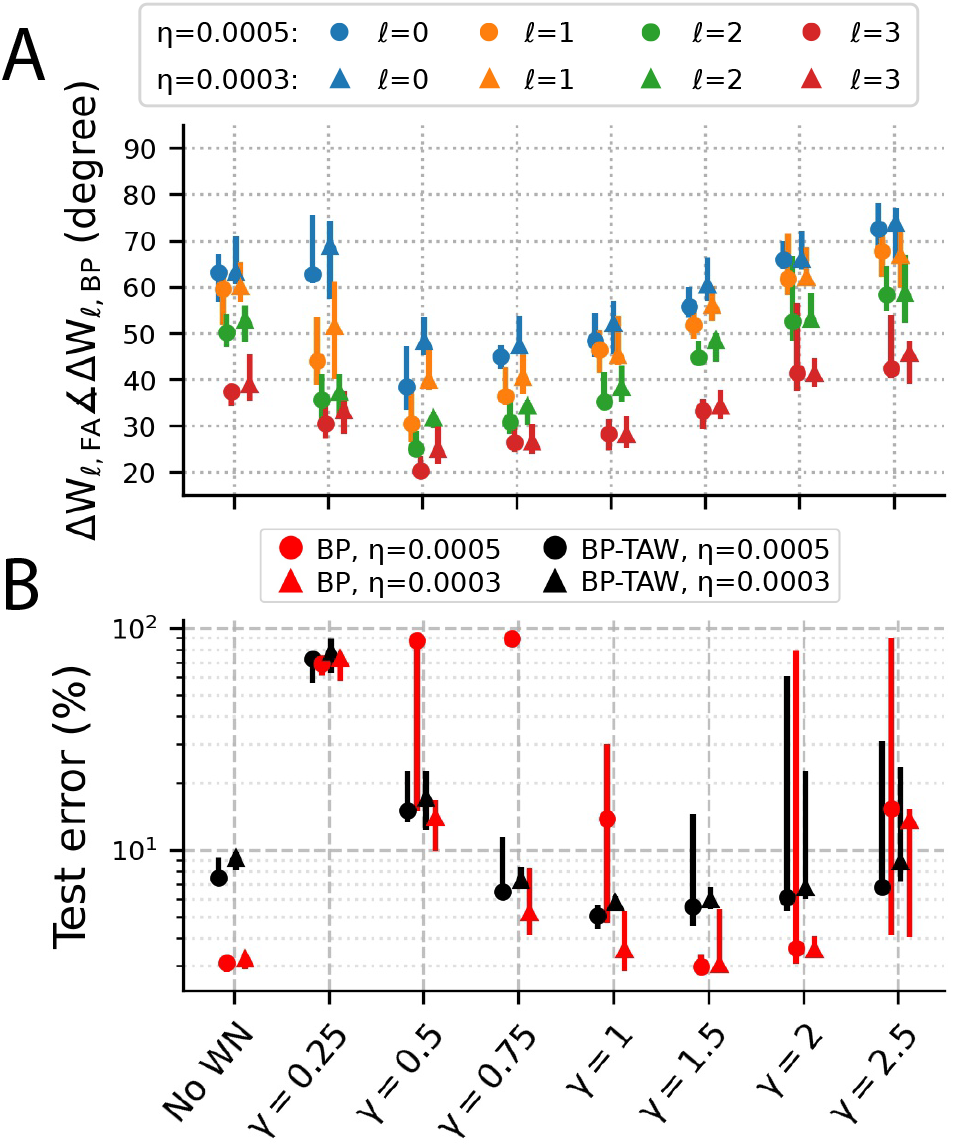
Sensitivity analysis and grid search for WN gain (*γ*) and learning rate (*η*) in the training of the example network on MNIST. **(A)** The angle between update directions of BP and those of BP-TAW for different values of WN gain (*γ*) separately for two learning rates *η* = 0.0003 and *η* = 0.0005 across all layers. **(B)** Classification error for different values of WN gain (*γ*) for BP and BP-TAW separately for two learning rates *η* = 0.0003 and *η* = 0.0005. (A,B) Each point is the median of 10 runs and error bars show range of the minimum and the maximum values.

### Supplementary Note 1| Taylor polynomials of higher degree

In the main text, we have obtained alignment terms by first order Taylor approximations. Therefore, each alignment term is the first-degree Taylor polynomial in an individual Taylor series. However, if more precision is required, and if the activation function is real and analytic, Taylor polynomials of higher degree can be expanded as follows:

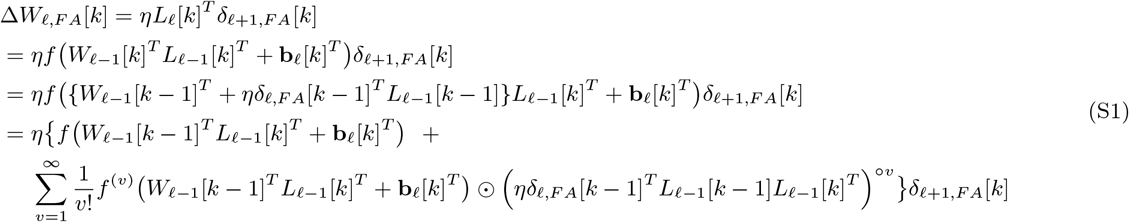

where (*·*)^*ov*^ denotes element-wise power of the matrix (also known as Hadamard power) and 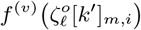 denotes the *n*-th derivative of the real analytic activation function *f* at the point 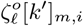. Analogously, we can expand *f(W*_𝓁 −1_[*k* − 1]^*T*^ *L*_𝓁 −1_[*k*]^*T*^ + **b**_𝓁_ [*k*]^*T*^) as follows

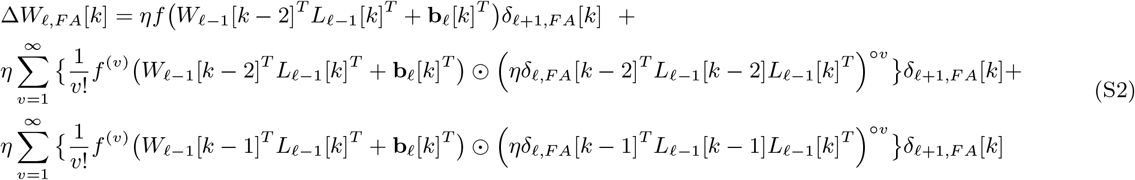

Doing this successively yields

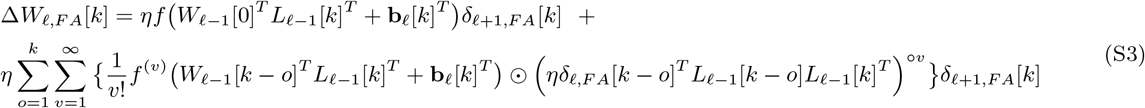

Taking 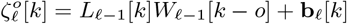 and *δ*_𝓁, *F A*_[*k* − *o*] = *δ*_𝓁 +1,*F A*_[*k* − *o*]*B*_𝓁_ ⊙ *f*^*′*^ (*Z*_𝓁_ [*k* − *o*]), Δ*W*_𝓁, *F A*_[*k*] reads

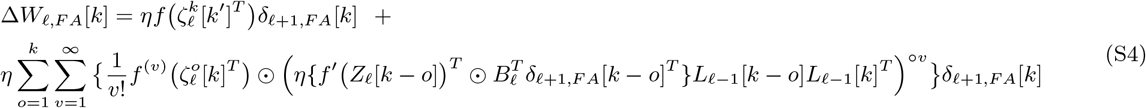

### Supplementary Note 2| Index notation of alignment terms

Index notation of alignment terms is very useful to gain further understanding about the underlying mechanism of FA and WA. According to the matrix notation of alignment terms that we have used in the main text, index notation of alignment terms can be driven as follows

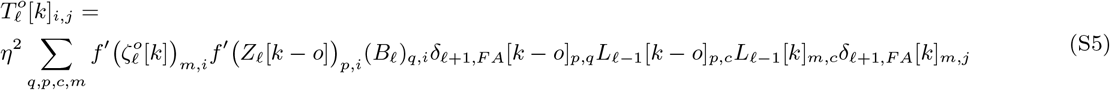

where the element in the *q*^*th*^ row and the *i*^*th*^ column of a matrix is denoted by a subscript consisted of a pair of small letters (for example (*B*_𝓁_)_*q,i*_ denotes the element in the *q*^*th*^ row and the *i*^*th*^ column of *B*_𝓁_).

Beyond individual alignment terms, we are interested in the behavior of the summation of all orders of alignment terms which appears in 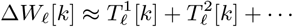. For a sample 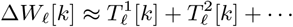 and a sample 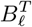, alignment in the sense of the injection of an aligned component into *W*_𝓁_ by the resultant of all orders of alignment terms at iteration *k*, namely,

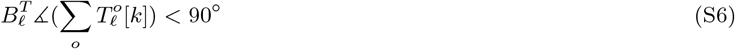

is equivalent to

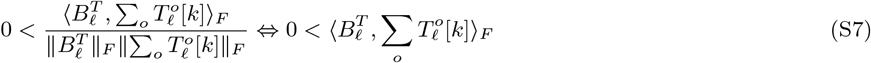

Therefore, the resultant of all orders of alignment terms at iteration *k* injects an aligned component into *W*_𝓁_ if and only if

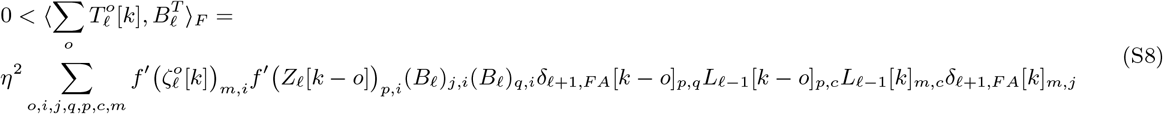

where 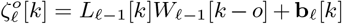 Equation S8 shows how input feedback weights to the layer 𝓁 ((*B*_𝓁_)_*j,i*_ and (*B*_𝓁_)_*q,i*_), error signals in the layer 𝓁 + 1 (*δ*_𝓁 +1,*FA*_[*k* − *o*]_*p,q*_ and *δ*_𝓁 +1,*FA*_[*k*]_*m,j*_), output signals in the layer 𝓁 − 1 (*L*_𝓁 −1_[*k* − *o*]_*p,c*_ and *L*_𝓁 −1_[*k*]_*m,c*_), and nonlinearity 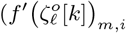 and *f ′(Z*_𝓁_ [*k* − *o*])_*p,i*_) are collectively contribute to alignment.

Considering the cosine similarity

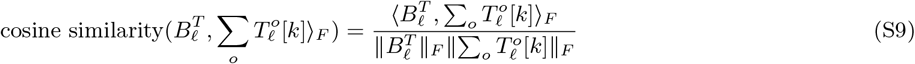

, since its denominator 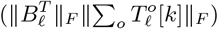 is nonnegative, if we want to only investigate occurrence of alignment regardless of its amount, we can simply refer to its numerator 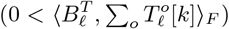 However, if we want to investigate its amount, we should also consider its denominator, especially 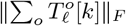we are less concerned about 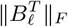 since in BP-TAW it is a fixed matrix across iterations).

The effect of nonlinearity as a distortion can be understood from the equation S8. If the activation function is an increasing function (which is a common choice in practical applications), nonlinearity scales individual terms in the summation of equation S8 with a nonnegative scalar and does not change their sign. In general, this can be a complex distortion which has some determinative and game-changing dependence with other signals. However, if this dependence is weak and nonlinearity does not have much contribution to the overall behavior of alignment terms, we can regard nonlinearity as a mild distortion and simply ignore it and refer to the linear case as follows

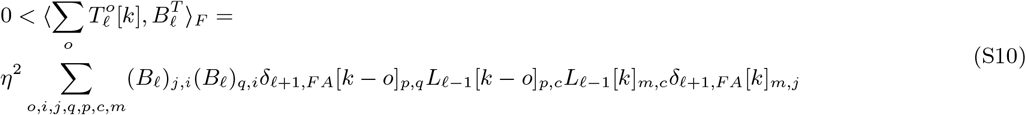

, otherwise we can still do our analysis by referring to the equation S8 and considering nonlinearity.

From the index notation of alignment terms, it can be understood that the process of BP-TAW sees mini-batches as the time sequence is folded in them. For example, shuffling data points within each mini-batch (not the whole data set as in the conventional data shuffling) is the same as changing the order of summations over *p* and *m* in the equations S8 and S5. It does not change the amount of 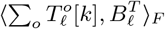 and also 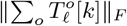; thus, it does not change the amount of alignment.

From the index notation of alignment terms, robustness of WA against data shuffling (shuffling the whole dataset at the beginning of each epoch) can be understood. Although data shuffling changes the trajectory of weight matrices, assuming the update steps to be small, in terms of statistical properties contributing to WA, shuffling is approximately similar to changing the order of summations over *o* and *p* in the equations S8 and S5. For example, from the point of view of the *k*^*th*^ mini-batch, a data point of the dataset that without data shuffling has appeared at the second data point (*p* = 2) of the (*k* − 20)^*th*^ mini-batch (*o* = 20), after data shuffling may appear at the tenth data point of the (*k* − 5)^*th*^ mini-batch (*o* = 5 and *p* = 10). If neural activities produced in the network by this data point in both of these positions are similar to each other, changing its position among lags can change the behavior of each alignment term of order *o* = 5 and *o* = 20 but does not change the behavior of their summation. Data shuffling does the same to all data points, but if the mentioned condition about the similarity of activity is approximately met, it approximately maintains the amount of 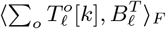and also 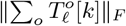and consequently the amount of alignment without data shuffling is approximately equal to the amount of alignment with data shuffling.

The contribution of autocorrelation of neural activity to the alignment can be understood from the index notation of alignment terms. For simplicity, consider the linear case with the assumption that three random vectors of [(*B*_𝓁_)_*j,i*_, (*B*_𝓁_)_*q,i*_], [*δ*_𝓁 +1,*F A*_[*k* − *o*]_*p,q*_, *δ*_𝓁 +1,*F A*_[*k*]_*m,j*_], and [*L*_𝓁 −1_[*k* − *o*]_*p,n*_, *L*_𝓁 −1_[*k*]_*m,n*_] are mutually independent, and also elements of *B*_𝓁_ are i.i.d. from *N* (0, *σ*^2^) (similar to the hypothetical condition of Fig. 2C corresponding to the blue trace). In this condition, the occurrence of alignment is expected if

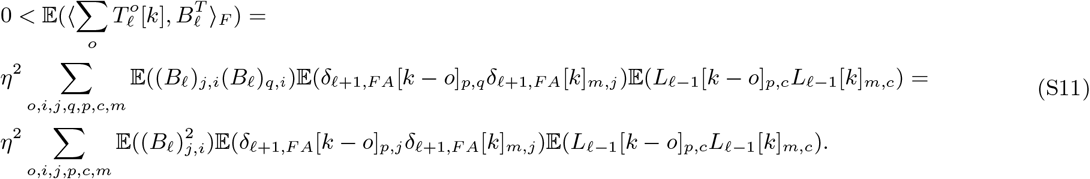

Accordingly, with the activation function chosen to be nonnegative (analogous to firing rate of biological neurons), alignment happens if the error signals (*δ*_𝓁 +1,*F A*_[*k* − *o*]_*p,j*_) and the outputs of neurons (*L*_𝓁 −1_[*k*]_*m,c*_) are positively autocorrelated among lags (summation on *o*) or mini-batches (summation on *m* and *p*), since as previously mentioned, the process of BP-TAW sees mini-batches as the time sequence folded in them. In other words, alignment happens if elements of *L*_0_ and *δ*_2_ are positively autocorrelated in the sense that

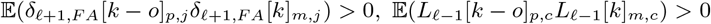

which is in accordance with the standard definition of autocorrelation function of a stochastic process. Note that in this hypothetical condition, which is proposed to provide intuition about WA, error and input signals are mutually independent. In general, these conditions are not necessarily met and in addition to autocorrelation, other properties of neural activity, such as cross-correlation between error and output signals, contribute to alignment.

The contribution of cross-correlation between error and output signals of neurons to the alignment can be understood from the index notation of alignment terms. For simplicity, consider the linear case with the assumption that three random vectors of [(*B*_𝓁_)_*j,i*_, (*B*_𝓁_)_*q,i*_], [*δ*_𝓁 +1,*F A*_[*k* − *o*]_*p,q*_, *L*_𝓁 −1_[*k* − *o*]_*p,n*_], and [*δ*_𝓁 +1,*F A*_[*k*]_*m,j*_, *L*_𝓁 −1_[*k*]_*m,n*_] are mutually independent, and elements of *B*_𝓁_ are i.i.d. from 𝒩 (0, *σ*^2^), and output and error signals are unautocorrelated in the sense that

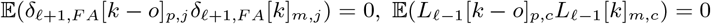

for *o* ≠ 0 (similar to the hypothetical condition of Fig. 2C corresponding to the green trace). In this condition the occurrence of alignment is expected if

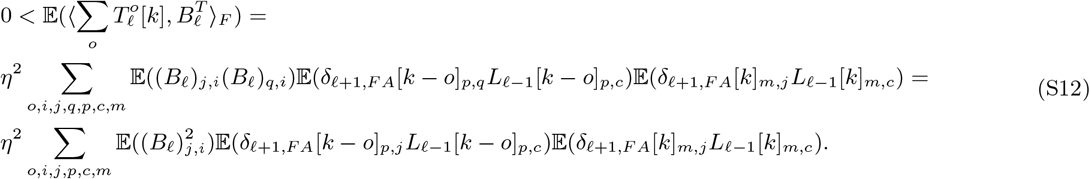

In other words, alignment happens if elements of *L*_0_ and *δ*_2_ are positively cross-correlated in the sense that

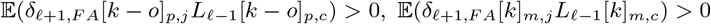

which in accordance with the standard definition of cross-correlation function of two stochastic processes. Note that in this hypothetical condition, which is proposed to provide intuition about WA, samples of error and output signals are independent among lags. In general, these conditions are not necessarily met.

All simplifying assumptions in this work are made to facilitate the analysis and explain the occurrence of alignment, whose underlying mechanism had remained elusive, in a simple and intuitive way. These assumptions do not weaken the generality of the provided framework because one can always refer to the nonlinear form of alignment terms (or even higher-order Taylor approximation) and also consider dependencies between backward weights and other signals, leading to a more elaborate analysis at the expense of simplicity.

### Supplementary Note 3| Skew-symmetric part of the transformation matrix totally deviates the direction

The skew-symmetric part of the transformation matrix (or any real skew-symmetric matrix) totally deviates *B*^*T*^ (or any real matrix) after matrix multiplication since

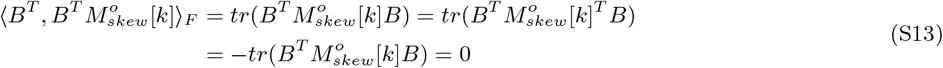

where *tr*(*·*) denotes matrix trace, and as a result 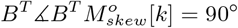. Hence, the amount of alignment is dependent on 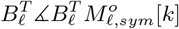 and the ratio of 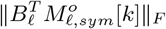 to 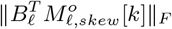.

### Supplementary Note 4| Direct feedback alignment

In DFA (Nøkland, 2016), error signals are directly backpropagated from the output layer to each hidden layer through direct fixed random backward weights denoted by 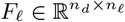as follows

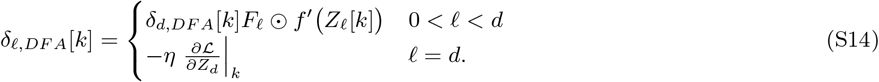

For the update directions of the backpropagation through direct random weights we can write

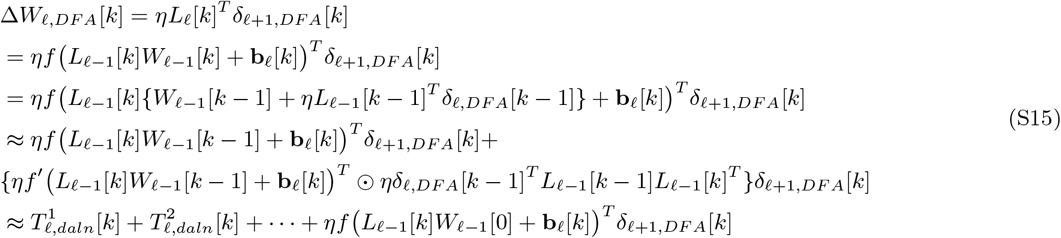

where we define *direct alignment term* of order *o* as bellow

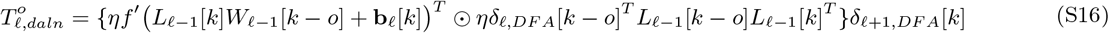

We can start from 𝓁 = *d* − 1 towards 𝓁 = 1 and decompose every 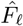such that for 𝓁 = *d* − 1 we have,

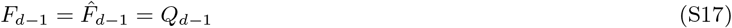

, and for 0 ≤ *𝓁 < d* − 1 we have

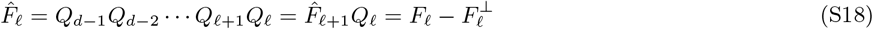

where *Q*_𝓁_ is the unique least squares solution of arg min 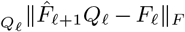 with minimum possible ║ *Q*_*d*_ *║* _*F*_ and 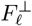is perpendicular to 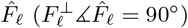. According to this decomposition, we can write

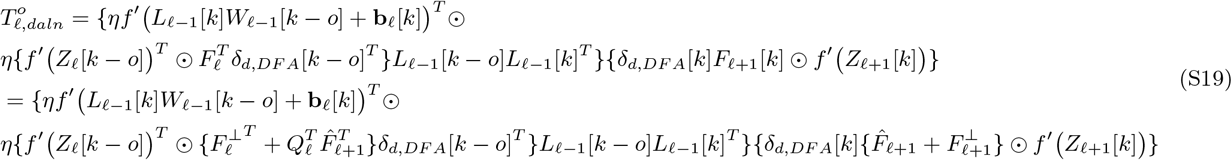

In the linear case, and by assuming 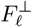 to be negligible (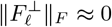 and 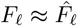, this can be a good assumption depending on the structure and dimension of direct backward weight matrices and the column space of 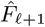), direct alignment term of order *o* is

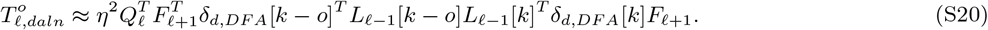

According to the structure of direct alignment terms in this simplified condition and with an analysis similar to the analysis of alignment terms in FA, the alignment of each *W*_𝓁_ with 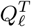 and consequently the alignment of 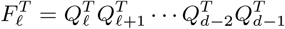 with *W*_𝓁_ *W*_𝓁 +1_ *· · · W*_*d*−2_*W*_*d*−1_ can be understood. We leave further investigation of DFA and its general condition to future works.

